# Integrative transcriptomic analysis of SLE reveals IFN-driven cross-talk between immune cells

**DOI:** 10.1101/2020.04.27.065227

**Authors:** Bharat Panwar, Benjamin J. Schmiedel, Shu Liang, Brandie White, Enrique Rodriguez, Kenneth Kalunian, Andrew J. McKnight, Rachel Soloff, Gregory Seumois, Pandurangan Vijayanand, Ferhat Ay

## Abstract

The systemic lupus erythematosus (SLE) is an incurable autoimmune disease disproportionately affecting women and may lead to damage in multiple different organs. The marked heterogeneity in its clinical manifestations is a major obstacle in finding targeted treatments and involvement of multiple immune cell types further increases this complexity. Thus, identifying molecular subtypes that best correlate with disease heterogeneity and severity as well as deducing molecular cross-talk among major immune cell types that lead to disease progression are critical steps in the development of more informed therapies for SLE. Here we profile and analyze gene expression of six major circulating immune cell types from patients with well-characterized SLE (classical monocytes (n=64), T cells (n=24), neutrophils (n=24), B cells (n=20), conventional (n=20) and plasmacytoid (n=22) dendritic cells) and from healthy control subjects. Our results show that the interferon (IFN) response signature was the major molecular feature that classified SLE patients into two distinct groups: IFN-signature negative (IFNneg) and positive (IFNpos). We show that the gene expression signature of IFN response was consistent (i) across all immune cell types, (ii) all single cells profiled from three IFNpos donors using single-cell RNA-seq, and (iii) longitudinal samples of the same patient. For a better understanding of molecular differences of IFNpos versus IFNneg patients, we combined differential gene expression analysis with differential Weighted Gene Co-expression Network Analysis (WGCNA), which revealed a relatively small list of genes from classical monocytes including two known immune modulators, one the target of an approved therapeutic for SLE *(TNFSF13B/BAFF:* belimumab) and one itself a therapeutic for Rheumatoid Arthritis *(IL1RN:* anakinra). For a more integrative understanding of the cross-talk among different cell types and to identify potentially novel gene or pathway connections, we also developed a novel gene co-expression analysis method for joint analysis of multiple cell types named integrated WGNCA (iWGCNA). This method revealed an interesting cross-talk between T and B cells highlighted by a significant enrichment in the expression of known markers of T follicular helper cells (Tfh), which also correlate with disease severity in the context of IFNpos patients. Interestingly, higher expression of *BAFF* from all myeloid cells also shows a strong correlation with enrichment in the expression of genes in T cells that may mark circulating Tfh cells or related memory cell populations. These cell types have been shown to promote B cell class-switching and antibody production, which are well-characterized in SLE patients. In summary, we generated a large-scale gene expression dataset from sorted immune cell populations and present a novel computational approach to analyze such data in an integrative fashion in the context of an autoimmune disease. Our results reveal the power of a hypothesis-free and data-driven approach to discover drug targets and reveal novel cross-talk among multiple immune cell types specific to a subset of SLE patients. This approach is immediately useful for studying autoimmune diseases and is applicable in other contexts where gene expression profiling is possible from multiple cell types within the same tissue compartment.

## INTRODUCTION

Systemic lupus erythematosus (SLE) is a chronic autoimmune disease that affects multiple organs including the skin, joints, the central nervous system and the kidneys^1,2^. It is caused by an aberrant autoimmune response to produce antibodies against self-antigens and these autoantibodies form immune complexes (ICs), which are then deposited into different organs and affect their normal function^3^. SLE is a highly heterogeneous disease in terms of development, presentation, manifestations, and severity; also the incidence and prevalence vary remarkably^4^. Thus, the time-course of flares and remission is unpredictable^5^. The diverse clinical manifestations of SLE present a challenge due to the involvement of several organs as well as diversified autoantibodies^6^. Therefore, accurate diagnosis and disease activity assessment is essential for managing SLE disease^7^. Even though the SLEDAI (SLE Disease Activity Index) score, which measures disease activity from 24 different clinical variables^8^, is widely adopted, it still leaves out several manifestations due to heterogeneous nature of SLE^9^. Such heterogeneity in manifestations and in assessing disease severity and activity has made it difficult to develop efficient therapeutics and, to date, only one drug, belimumab, which targets the *TNFSF13B* gene encoding the B-cell activating factor, BAFF, has been approved for use in SLE^1^.

Clinical heterogeneity is the main obstacle in finding good treatments for SLE^10^, therefore it is important to discover molecular subtypes and signatures that correlate to clinical phenotypes. One strong signature characterized to date is the heightened expression levels of type I interferon (IFN) related genes in blood transcriptome of SLE patients and in correlation with disease activity^11–13^ and pathogenesis^5^. Additionally, a recent single-cell transcriptome study revealed the upregulation of IFN-inducible genes in Lupus nephritis patients^14^. However, how this IFN response affects molecular programs of different immune cells and how it influences the cross-talk among different types of immune cells is largely unknown as most studies use mixed populations of immune cells (e.g., PBMCs). Therefore, transcriptome profiling with sorted populations of major immune cell types provides an opportunity to define cell-specific molecular programs and correlate signatures between cell types. The traditional approach is to study each cell type one-by-one and to find differentially expressed genes (DEGs) for those cells based on the gene-level expression estimates within and across different groups. This approach is complemented by analyzing the patterns of correlated gene expression (co-expression) which relies on the “guilt-by-association” principle^15^, suggesting genes with coordinated changes in expression are more likely to be involved in similar biological functions. Differential network analysis uses these co-expression patterns across different conditions to identify differentially connected genes (DCGs), revealing irregularities in the transcriptome wiring in the disease state 16-20

The major goals of this study are to advance our understanding of the clinical and molecular heterogeneity in SLE and to have a systematic view of the molecular cross-talk among different immune cells that collectively contribute to disease pathogenesis. In order to achieve these objectives, we have generated gene expression profiles of six major human immune cell types from a cohort of SLE patients and matched healthy controls using RNA-seq^21^. Our cohort consists of samples from 64 SLE patients (62F, 2M) and 24 controls (all female) as well as their demographic information, clinical features (e.g., SLEDAI) and measurements of the plasma levels of several relevant cytokines and chemokines. From the gene expression profiles of different immune cells, we found that the IFN response is a determining factor in the stratification of SLE patients into two distinct groups independent of which cell type is considered. We performed differential expression and differential network analysis between IFN-positive (IFNpos) and IFN-negative (IFNneg) patients to prioritize a gene list that likely drives the generally more severe IFNpos phenotype. This analysis led to a relatively small list of genes from classical monocytes including two known immune modulators, one the target of an approved therapeutic for SLE *(TNFSF13B/BAFF: belimumab)* and one itself a therapeutic for Rheumatoid Arthritis *(IL1RN: anakinra).*

As multiple immune cell types are involved in immuno-pathogenesis of SLE and they likely act in coordination^22,23^, which renders the approach to studying each cell type in isolation suboptimal. The standard co-expression network analysis tool ‘WGCNA’ clusters the well coexpressed genes into discrete modules^24^, for one cell type at a time. In order to coordinate the study of all six cell types we profiled, we developed a novel integrated WGCNA (iWGCNA) approach that extends the WGCNA framework. Although we primarily focus on SLE in this work, the iWGCNA framework can be applied to any context where multiple immune cell types coexist to explore cross-talk at the gene and the pathway level *in silico*. When applied to data from classical monocytes (cMo), B cells, T cells, neutrophils (PMN), and conventional (cDC) as well as plasmacytoid dendritic cells (pDC), iWGCNA also confirmed the dominance of the IFN signal in stratifying the SLE patients by clustering IFN-related genes from different cell types into one particular module. The iWGCNA approach also reported an interesting module that reveals a potential cross-talk between T and B cells highlighted by a significant enrichment in the expression of known markers of T follicular helper cells (Tfh), overexpression of which also correlates with disease severity for the IFNpos SLE patients. The expansion of circulating T cells resembling Tfh also has been shown in mouse models of SLE as well as subgroup of patients with SLE^25^. However, the connection between Tfh cells and the interferon signature is a new discovery to the best of our knowledge. Tfh cells in the germinal centers as well as in circulation (cTfh)^26^ are known to promote B cell class-switching and help the development of high affinity antibodies^27^. More recently, a study described an expanded population of CD4^+^ helper T cells in SLE patient blood that are distinct from Tfh but still help activate B cells, further confirming heightened cross-talk between circulating B cells and T cells in SLE^28^.

An important factor in the context of B cell help is, *BAFF*, which promotes B cell survival and controls their maturation^29^ while also regulating IFN-gamma production by Tfh cells in lupus-prone mice^30^. Increased expression of *BAFF* induces expansion of activated B-cells that produce auto-antibodies and cause autoimmunity and is dependent, at least in part, on Tfh^31^. Interestingly, in our data, higher expression of *BAFF* from all myeloid cells shows a strong correlation with the enrichment of the Tfh gene signature in T cells, which itself is elevated in IFNpos SLE patients. Therefore, our novel analysis links the IFN response signature shared by all immune cell types to higher expression of *BAFF* in myeloid cells and Tfh related genes in T cells, and to heightened B cell help leading to robust antibody production. We anticipate this data-driven and integrated analysis of multiple immune cell types, which allowed us to prioritize known modulators of immuno-pathogenesis and discover interesting connections across multiple cell types in SLE, will also be immediately useful for expanding our understanding of SLE and other autoimmune diseases.

## RESULTS

### Transcriptional profile of classical monocytes reveals two molecular subtypes of SLE

Since SLE is known for its heterogeneity, we first investigated the full set of patients in our cohort for distinct molecular signatures. To this end, we analyzed the bulk RNA-seq based transcriptome profiles of classical monocytes (cMo) from 64 SLE patients and compared them to that of 24 healthy controls (HC) (**Figure 1A; Supplementary Figure S1A-B**). For this analysis we only used the samples gathered from the first study visit of each patient, setting aside the longitudinal samples for validation. This analysis revealed a total of 125 differentially expressed genes (DEGs; P.adj < 0.05) with 109 up- and 16 down-regulated genes. The PCA plot of these 125 DEGs highlights the heterogeneity within SLE patients, with one group showing clear differences compared to HCs and the other showing no such separation (first principal component; **Figure 1B**). Further analysis of these 125 DEGs showed that most of these DEGs are enriched in interferon (IFN) related pathways (**Figure 1C**; **Supplementary Figure S1C-D**, details in **Supplementary Table S1**). For further illustrating this point, we used 20 interferon signature genes (IFN-20) for Gene Set Enrichment Analysis (GSEA;^32^) and showed a significant enrichment of their expression in SLE compared to HC (Normalized Enrichment Score (NES) = 0.962; FDR q-value = 0.023; **Figure 1C**). We also used the IFN-20 set to further classify SLE patients into two subgroups (IFNpos and IFNneg) with clear differences in their molecular signatures (**Figure 1D, Supplementary Figure S1D**) including the longitudinal samples (**Supplementary Figure S1E**). To better quantify these differences, we conducted pairwise comparisons of the resulting three groups (IFNpos, IFNneg, HC) identifying 1439 DEGs between IFNpos and IFNneg and 1288 DEGs between IFNpos and HC, whereas there were only 5 DEGs between IFNneg and HC (P.adj < 0.05; **Supplementary Table S1**). These results suggest that it is important to investigate these two molecular sub-groups of SLE (IFNpos and IFNneg) separately and that there is no clear cMo molecular signature that distinguishes IFNneg patients from HC. This distinction is likely also critical for developing therapeutics for SLE as their clinical testing may suffer from such high heterogeneity among the patient cohort.

**Figure 1.**
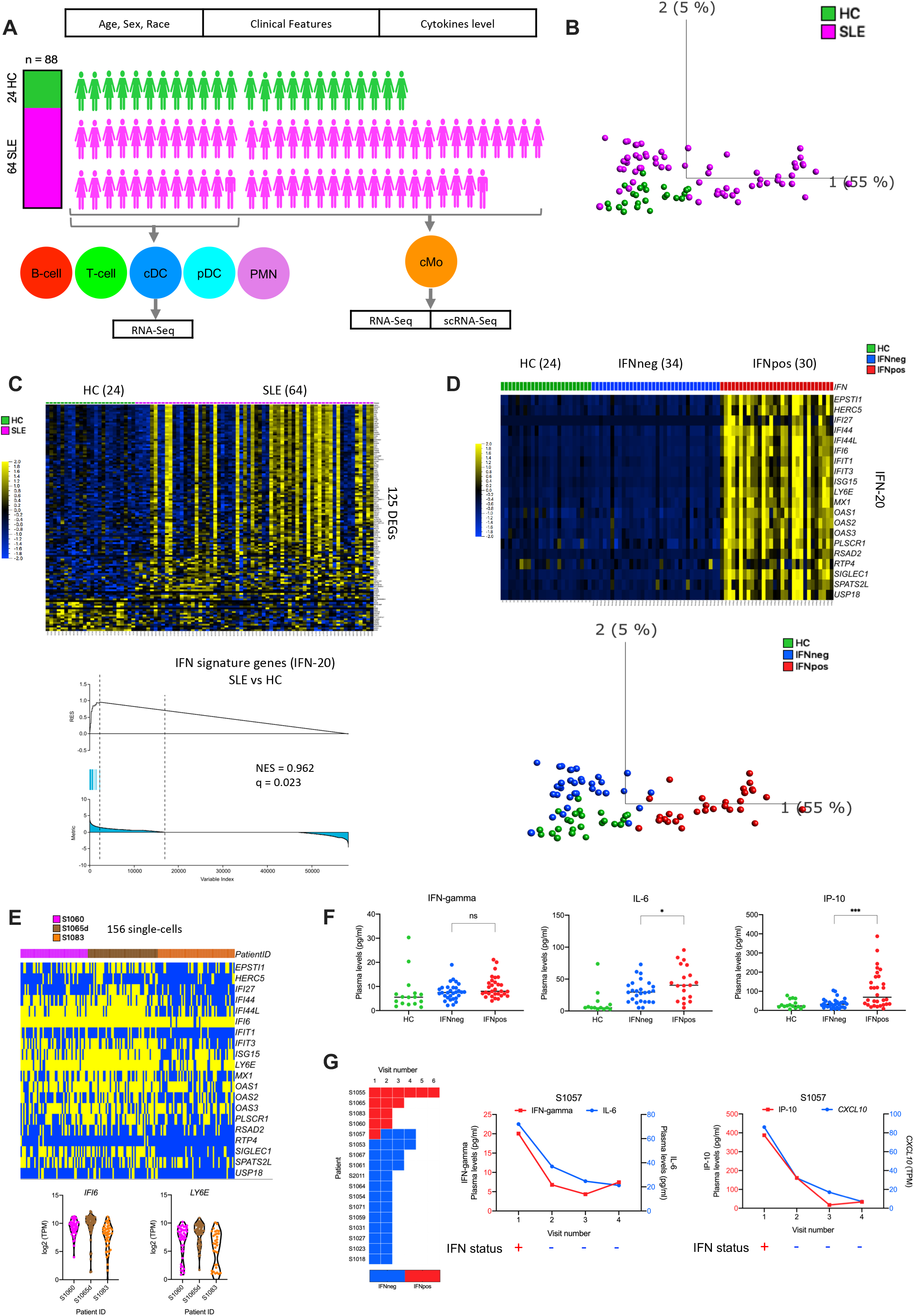
Transcription profile of classical monocytes reveals two molecular subtypes of SLE that shows stability in longitudinal data. **(A)** Overview of the SLE cohort. Healthy control (HC) samples are highlighted in green color and SLE sample are highlighted in magenta color. Six different immune cell types including classical monocytes (cMo), Polymorphonuclear Neutrophils (PMNs), conventional dendritic cells (cDC), plasmacytoid dendritic cells (pDC), T-cells, and B-cells are displayed in different colors. **(B)** The PCA plot based on 125 DEGs (P.adj < 0.05 from Benjamini-Hochberg test in DESeq2) between SLE and HC in classical monocytes. Green and magenta colors are representing HC and SLE patients, respectively. **(C)** Heatmap of DEGs (one per row) in SLE versus HC (P.adj < 0.05) comparison and presented as row-wise z-scores of transcripts per million (TPM) in SLE (magenta) or HC (green); each column represents an individual patient. Only first study visit data were used for differential analysis and the number of individual samples is noted in brackets. The GSEA plot shows significant enrichment (NES=0.962; q=0.023) of 20 IFN-signature genes (IFN-20) in SLE vs HC. **(D)** SLE patients (first study visits) classified into two groups based on the expression of IFN-20 genes. Where each gene is presented as row-wise z-scores of transcripts per million (TPM) in IFNpos (red), IFNneg (blue) and HC (green); each column represents an individual patient. The PCA plot shows two molecular sub-types of SLE in different colors. Green, blue, and red colors are representing HC, IFNneg, and IFNpos respectively (we used this color scheme in all figures). **(E)** The binary heatmap plot of single cell level expression of IFN-20 in cMo (156 cells) from three different IFNpos patients (shown in different colors in columns). In this binary heatmap, expression level of >1 TPM is highlighted in yellow and <1 TPM is shown in blue. The expression of two genes (*IFI6* and *LY6E*) is also provided as violin plots. **(F)** Scatter dot-plots show the expression level (pg/ml) in plasma of IFN-gamma, IL-6 and IP-10. Only first study visit samples (n=76) were used in these scatter dot-plots. The difference between IFNpos and IFNneg has been calculated using unpaired T-test (two-tailed) and the statistical significance (p-value) level is shown in the plots (ns: not significant, *: <0.05, ***: <0.001. **(G)** Longitudinal IFN response status of 17 SLE patients (5 patients were IFNpos and 12 patients were IFNneg at the first study visit) (left). Only one patient (S1057) changed their IFN response status between study visits, changing from IFNpos to IFNneg between the first and second study visit and remained IFNneg in all follow-up visits. Two plots with connecting lines (middle and right) show expression changes of multiple cytokines and chemokines in the longitudinal data for patient S1057, where different analytes are plotted on different Y-axes (IFN-gamma on Y1-axis in red color and IL-6 on Y2-axis in blue color) in different colors. The RNA-level expression of *CXCL10* is also shown (Y2-axis) with protein-level expression of IP-10 (Y1-axis). The lower panel of each plot shows their IFN response status over multiple longitudinal visits.

The bulk RNA-seq data provides average gene expression but lacks the resolution to determine the proportion of cMo that are positive for the expression of IFN signature genes. To this end, we used single-cell RNA-seq to test whether the IFNpos status is conferred by a subset of cells or is shared across each individual cMo cell. Our analysis of the transcriptome profiles of ~156 single cells from 3 different IFNpos patients clearly demonstrates that nearly all cMo are positive for the expression of at least several of the IFN-20 genes (**Figure 1E**). Specially, *IFI6* and *LY6E,* both well-known type I interferon related genes^33^, are highly expressed in most cMo across different patients (violin plot; **Figure 1E**). These results suggest that the IFN response signature is not specific to a subset of classical monocytes but is a robust feature across the cell population.

### IFN response status of SLE patients correlates with the levels of pro-inflammatory cytokines and chemokines

For a subset of donors from our cohort, we measured the plasma level of different cytokines and chemokines such as IFN-alpha, IFN-beta, IFN-gamma, IL-2, IL-6, IL-10, IP-10, TNF-alpha, and TGF-beta and compared the levels from only the first study visit data across the three groups: HC (n=16), IFNneg (n=30) and IFNpos(n=30). Out of these IFN-alpha (p=0.0190), IFN-beta (p=0.0460), IL-6 (p=0.0413), and IP-10 (p=0.0002) levels were higher in SLE subgroups in comparison to HC. These three cytokines (IFN-alpha, IFN-beta, IL-6) as well as IP-10 had a statistically significant difference between IFNpos and IFNneg SLE patients as well (**Figure 1F; Supplementary Figure S1F**). The role of IFNs as pro-inflammatory cytokines is well-studied in SLE^34,35^. Another cytokine with a significant difference between IFNpos and IFNneg is IL-6, which is also pro-inflammatory. IL-6 induces the maturation of B-lymphocytes into plasma cells and increases Ig secretion. Increasing IL-6 levels correlate with increased disease activity and with anti-DNA autoantibody levels in human SLE^36^. It has also been shown that IL6-174G/C and IL6-572G/C polymorphisms are associated with the development of SLE^37^. We also identified a significantly higher level of a pro-inflammatory chemokine, interferon-inducible protein-10 (IP-10) in IFNpos patients (**Figure 1F**). The levels of IP-10 has been also reported to correlate with SLE disease activity and with organ manifestations of this disease^38^.

### IFN response status molecular signature is mainly conserved in longitudinal samples

Next, we asked whether the IFN response status, IFNneg or IFNpos as measured by the IFN-20 molecular signature, changes across different visits of the same patient (**Supplementary Figure S1E**). We observe that for the majority of the patients the IFN response status is identical across all their follow-up visits, regardless of changes in disease activity, flares and/or treatment regimens (e.g. prednisone use and dose) across the multiple visits. Among the 17 SLE patients with longitudinal data from multiple visits that are at least 1 month apart, 12 patients were IFNneg and 5 patients were IFNpos at their first study visit (**Figure 1G**). Out of these, only one patient (S1057) showed a change in the IFN response status with the first collection being IFNpos and the next three follow-up visits being IFNneg (**Figure 1G** – left panel). Interestingly, serum levels of IP-10 significantly dropped (2.4 fold) in line with switching from IFNpos to IFNneg between the first and second sample collections (**Figure 1G** – mid panel). This sharp decrease is also reflected as a 2.7-fold decrease in the mRNA levels of *CXCL10,* which encodes IP-10, in classical monocytes between the same two visits (**Figure 1G** – right panel). Although, the differences in IFN-gamma levels did not reach statistical significance (p=0.0644) when only first study visit data are compared between IFNpos and IFNneg (**Figure 1F**), for this specific patient, IFN-gamma levels also showed a 3-fold decrease in the second visit (**Figure 1G** - mid panel). Also, consistent with comparisons from the first study visit data, the level of IL-6 also showed a 2-fold decrease for the second visit of this patient (**Figure 1G** - mid panel). Though limited to a single patient with IFN response status change in our cohort, this observation highlights the importance of analyzing longitudinal samples in order to draw more robust conclusions from changes in cytokine and chemokine levels with respect to IFN gene signature, disease status and therapeutic interventions. Overall, these results show that IFN-based stratification of SLE patients is relatively stable across their longitudinal samples with changes in that gene expression-based status strongly reflected in changes in the plasma levels of specific pro-inflammatory cytokines and chemokines.

### Combined analysis of differential network and gene expression of classical monocytes reveals known immune modulators

The major limitation with differential gene expression analysis is that it treats each gene individually while comparing expression profiles, whereas most biological functions are performed by a group of genes working together in coordination. Therefore, we used weighted gene co-expression network analysis (WGCNA) to generate a network from classical monocyte gene expression profiles of 64 SLE donors^24^. This co-expression network revealed a total of 25 different modules, represented by different colors, where each module is a cluster of coexpressed genes. Furthermore, the correlation of eigengene value of each module with external clinical features showed that IFN response status is the most-correlated feature in comparison to other available clinical features (module-trait relationships) such as age, ethnicity, flare, severity, SLEDAI and treatment regimens (**Figure 2A**). The blue module is positively correlated (R = 0.83; P-value = 5e-17) with IFN response status (**Figure 2B**) and it consists of 1684 genes. In addition to our IFN-20 gene set, we also manually curated a broader set of 363 IFN-related genes (IFN-363) from different sources and found that most of the hub genes (genes with highest connectivity in the co-expression network) in the blue module are part of this IFN-363 set (**Supplementary Figure S2A**). The red module is negatively correlated (R = −0.73; P-value = 1e-11) with IFN response status and it has 662 genes, most of which are related to protein translation. This WGCNA analysis suggests that the dominance of IFN response signature in the stratification of SLE patients using gene expression is conserved when we use gene coexpression relationships instead.

**Figure 2.**
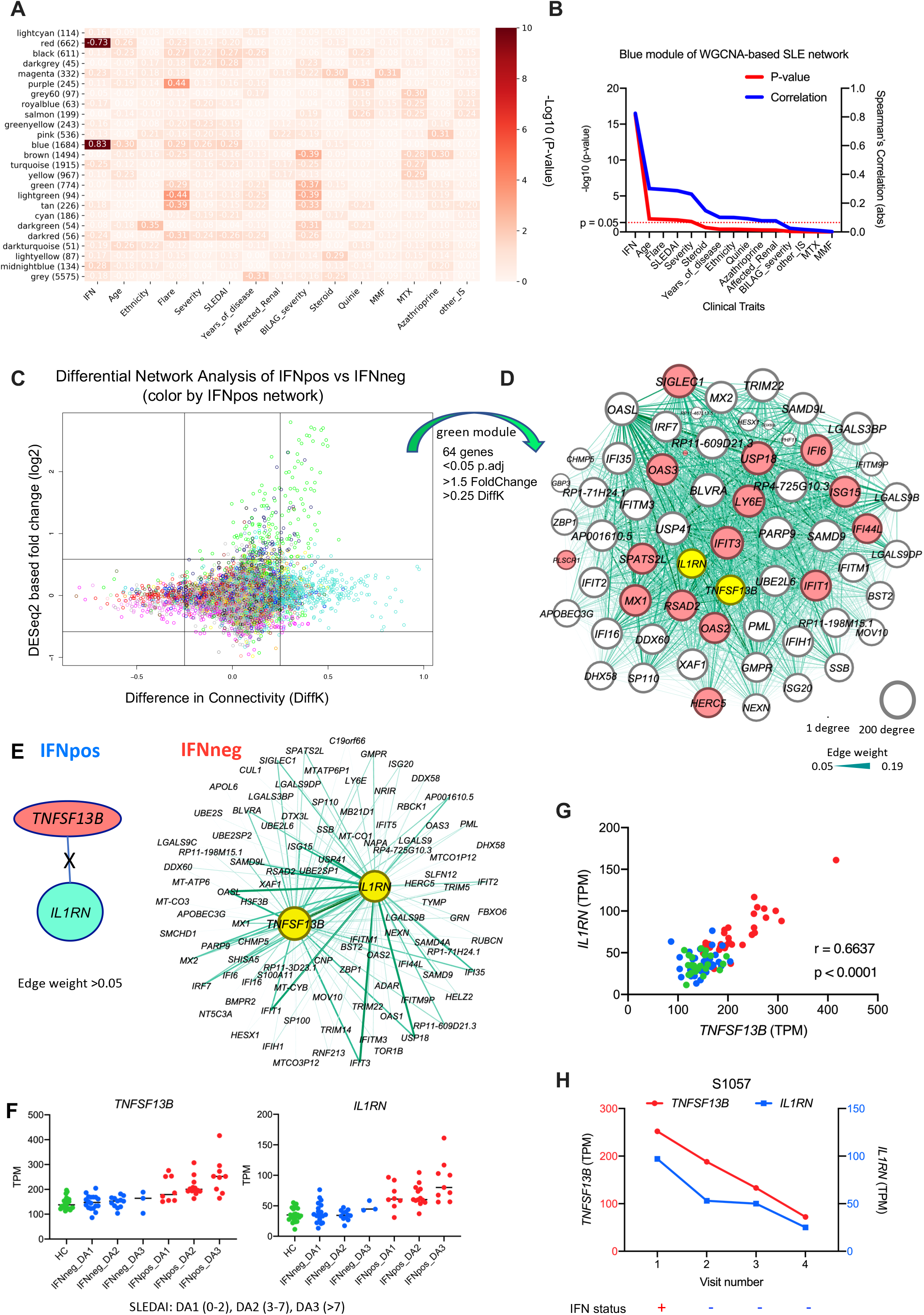
Combined analysis of differential network and gene expression of classical monocytes reveals two known immune modulators (*BAFF* and *IL1RN*) and shows their expression is dysregulated in SLE. **(A)** The module-traits correlation matrix heatmap of WGCNA based on total 25 different modules and external clinical features such as IFN, age, ethnicity, flare, severity etc. The color shows significance (-Log10 of p-value) and Spearman’s correlation values are provided in the heatmap. The number of genes in each module is also shown (in brackets) with the module name. IFN response status is the most correlated clinical feature and the blue module is the top positively correlated (r = 0.83; p = 5e-17) and the red module is the top negatively correlated (r = −0.73; p = 1e-11) with the IFN response status. **(B)** The line plot shows significance (-Log10 (P-value) on the Y1-axis in red color) and Spearman’s correlation (Y2-axis in blue color) of different clinical traits with eigengene values of the blue module. The clinical traits are sorted (decreasing order) based on their statistical significance (-Log10 of P-value) for the blue module. **(C)** The plot shows both differentially expressed (DEGs) and connected genes (DCGs) between IFNpos and IFNneg where the X-axis is the difference in the connectivity (DiffK = K1-K2; K1=connectivity in IFNpos network; K2=connectivity in IFNneg network) and the Y-axis is the DESeq2 based fold-change (log2). Colors of genes represent different modules based on the IFNpos (Network1) based network. **(D)** 64 genes were selected from the green module using significant DEGs (p.adj < 0.05; 1.5-fold change) as well as DCGs with at least 0.25 difference in connectivity (DiffK) and visualized by Gephi software. Where nodes are sized according to the number of edges (connections), and the edge thickness is proportional to the strength of co-expression. Available IFN signature genes (IFN-20) are highlighted in red colors and two known immune modulators (*IL1RN* and *BAFF* (*TNFSF13B*)) are highlighted in yellow. **(E)** An example showing the importance of differential network analysis because *IL1RN* and *BAFF*, which are in same green module, have a large number of connected genes in the IFNpos network whereas no connected genes (at threshold of 0.05 edge weight) are found in the IFNneg network. The strength of co-expression is also varying and is presented by the width of the connection. Both *BAFF* and *IL1RN* are in different modules (salmon and turquoise, respectively,) in the IFNneg network. **(F)** Expression of *BAFF* and *IL1RN* is plotted according to IFN response status and SLEDAI class. The SLEDAI score is divided into three different categories based on score: DA1 (0-2), DA2 (3-7), and DA3 (>7). Green, blue, and red colors represent HC, IFNneg and IFNpos, respectively. **(G)** The Spearman’s correlation (r = 0.6637; p < 0.0001) between *BAFF* and *IL1RN* based on RNA expression (TPM). **(H)** Expression changes of *BAFF* and *IL1RN* in longitudinal data from patient S1057, where genes are plotted on different Y-axes; *BAFF* on Y1-axis in red and *IL1RN* on Y2-axis in blue. The lower panel of this plot shows corresponding IFN response status over multiple longitudinal visits.

Though this WGCNA analysis on all SLE samples confirms our findings from the differential expression analysis, it does not help in prioritizing genes that may be potential therapeutic targets or are upstream regulators, leading to a large number of differentially expressed genes between the two groups. Therefore, we adopt a differential WGCNA approach, which has been useful in identifying differentially regulated genes^17^, to identify a set of differentially connected genes (DCGs) between IFNpos and IFNneg groups. Conceptually, a DCG may have the same level of expression in two different groups but may be co-expressed with distinct sets of genes suggesting a rewiring of the underlying biological pathways involving this gene. A gene that is both differentially expressed and differentially connected is also likely to have a broader impact in the molecular profile that distinguishes the compared groups from each other. With this motivation, we applied WGCNA to generate two separate co-expression networks, one for IFNpos and one for IFNneg, to identify DCGs between the two conditions. For each gene in each co-expression network, we computed network connectivity as the sum of connection strengths (based on co-expression) with the other genes in that network. We identify DCGs by computing the difference between the connectivity (DiffK) values for each gene between IFNpos and IFNneg network. Using stringent criteria, we identified a total of 99 genes that are DEGs (1.5-fold change; Padj < 0.05) as well as DCGs (> 0.25 or < −0.25 DiffK) when IFNpos and IFNneg patients are compared. A majority of these genes (64/99) are from the green module which includes known IFN related genes as well as several other genes previously implicated in the context of SLE (**Figure 2C, Supplementary Table S2**). A joint visualization of differential expression and connectivity highlights that the green module genes are enriched in significant differences in both expression as well as connectivity (**Supplementary Figure S2B**) with enrichment in IFN-related pathways (**Supplementary Figure S2C**). Interestingly, aside from 16 genes from IFN-20 set, the green module also harbors several hub genes such as *TNFSF13B, IL1RN, BLVRA, ZBP1, PARP9, APOBEC3G, LGALS9DP, GMPR,* and *USP41* (**Figure 2D**). Strikingly, this short list of genes includes two known immune modulators, *BAFF (TNFSF13B)* and *IL1RN* (highlighted in **Figure 2D**). *TNFSF13B,* also known as *BAFF* (B-cell activating factor), is the target of the only approved therapeutic (belimumab) for SLE to date^1^. A recombinant form (Anakinra) of the other gene, *IL1RN* (interleukin-1 receptor antagonist), is a therapeutic agent marketed for the treatment of Rheumatoid Arthritis^39^ and it inhibits the binding of pro-inflammatory IL1-α and IL1-β to the IL1 receptor^40^.

### *BAFF* and *IL1RN* expression is dysregulated in SLE

*BAFF* is significantly differentially expressed (P.adj = 8.14E-16; 1.55-fold change) as well as differentially connected (DiffK = 0.401) between IFNpos and IFNneg patients (**Supplementary Table S2**). *IL1RN* is also significantly differentially expressed (P.adj = 3.20E-11; 1.8-fold change) and differentially connected (DiffK = 0.330). No genes are connected to *BAFF* or *IL1RN* in IFNneg patients (edge weight > 0.05). In contrast both genes are hubs in the green module of the IFNpos network (**Figure 2E**). The expression levels of both *BAFF* and *IL1RN* show a strong positive correlation with SLEDAI score for IFNpos patients (**Figure 2F**). This correlation is valid when the continuous SLEDAI score is used or when this score is divided into three different disease activity categories: DA1 (0-2), DA2 (3-7), and DA3 (>7) as previously described^34^. The expression levels of *BAFF* and *IL1RN* are also strongly correlated with each other (r = 0.6637; p < 0.0001; **Figure 2G**). In the longitudinal data, the expression levels of both genes go down when the IFN response status of patient S1057 changes from IFNpos to IFNneg (from the first to second collection; **Figure 2H**). The DICE database,^41^, which profiled gene expression of nearly 100 healthy donors for 13 distinct human primary immune cell populations (including cMo, T and B cells), also shows that the expression of *BAFF* and *IL1RN* are specifically high in classical monocytes in the circulating blood (**Supplementary Figure S2D)**.

Additionally, this differential expression and connectivity analysis also revealed many interesting novel genes that have similar expression patterns to *BAFF* and *IL1RN* including the interferon-related genes *IFI6, LY6E, IFIT3* (**Supplementary Figure S2E**). For example, *BLVRA* (biliverdin reductase A) has a canonical function to convert biliverdin to bilirubin but it plays an anti-inflammatory function by activation of the PI3K–Akt-IL-10 pathway as well as inhibition of *TLR4* expression via direct binding to the *TLR4* promoter^42^. *ZBP1* (Z-DNA-binding protein 1) is a cytosolic DNA sensor that can activate type I IFN response^43^ and it also regulates programmed cell death and other inflammatory responses^44^. *DDX60L* (DExD/H-Box 60 Like) is an IFN-stimulated gene and is involved in anti-viral immunity^45^. *APOBEC3G* (Apolipoprotein B MRNA Editing Enzyme Catalytic Subunit 3G) is related to RNA editing and also plays an important role in anti-viral immunity^46^. *GMPR* (Guanosine Monophosphate Reductase) was recently reported as a potential therapeutic target for Alzheimer’s disease^47^. The revelation of these validated immune-related genes shows the robustness of our data-driven and unbiased bioinformatics analysis in identifying known and novel therapeutic targets for SLE and potentially for other autoimmune diseases, in general.

### IFN response molecular signature is conserved across multiple different immune cell types

The recent studies highlight that multiple immune cell types are involved in the pathogenesis of SLE^23^. Therefore, it is important to systematically investigate distinct immune cell types from the same cohort of patients and to jointly study them in the context of disease heterogeneity and disease severity. Here we used RNA from FACS-sorted populations of six different immune cell types including classical monocytes (cMo), polymorphonuclear neutrophils (PMNs), conventional dendritic cells (cDC), plasmacytoid dendritic cells (pDC), total T cells, and total B cells (**Supplementary Figure 1B**) and perform bulk RNA-seq to understand the stability of IFN response signature as well as the importance of cell type-specific features in SLE pathogenesis. For this multi-cell type profiling, we use a matched set of HCs, IFNneg and IFNpos samples (n=9-12 for each) from the first sample collections of different donors. The principal component analysis of these different cell types across each donor based on the top-1000 most variable genes reveals that each cell type clusters separately as expected (**Figure 3A**). We then carried out PCA using only the expression of the 20 genes in the IFN-20 set and colored each sample either by its cell type (**Figure 3B** – left panel) or by the IFN response status of the donor, which we determined from cMo data as previously described (**Figure 3B** – right panel). This analysis highlights that for each cell type, the IFNpos samples (red) are clearly separated from IFNneg and HC samples (green and blue, respectively) by the first principal component (**Figure 3B**). The individual PCA plots for each cell type also show that the IFN-based annotations are cell type independent and can be assigned at the patient level (**Supplementary Figure S3A-F**). Heatmap visualization using IFN-20 gene set confirms this separation for each donor and for each cell type both when the 20 genes are aggregated (**Figure 3C**) and when visualized separately (**Supplementary Figure S3G**). Out of 12 IFNpos patients, 9 are classified into higher disease activity classes (DA2 or DA3) with no DA3 patients in the IFNneg group, suggesting that IFN-based molecular stratification of SLE patients is also clinically relevant (**Figure 3C**). Analysis of *BAFF* and *IL1RN* gene expression, originally identified from our cMo analysis, show that both are expressed at high levels in PMN cells and are specifically upregulated in IFNpos samples (**Figure 3D**), likely contributing to disease activity. Together with the results discussed before, we established that our classification with respect to IFN response status correlates with disease activity and is robust to the mode of gene expression profiling (bulk or single-cell) and to the immune cell type under investigation, supporting its clinical relevance in SLE.

**Figure 3.**
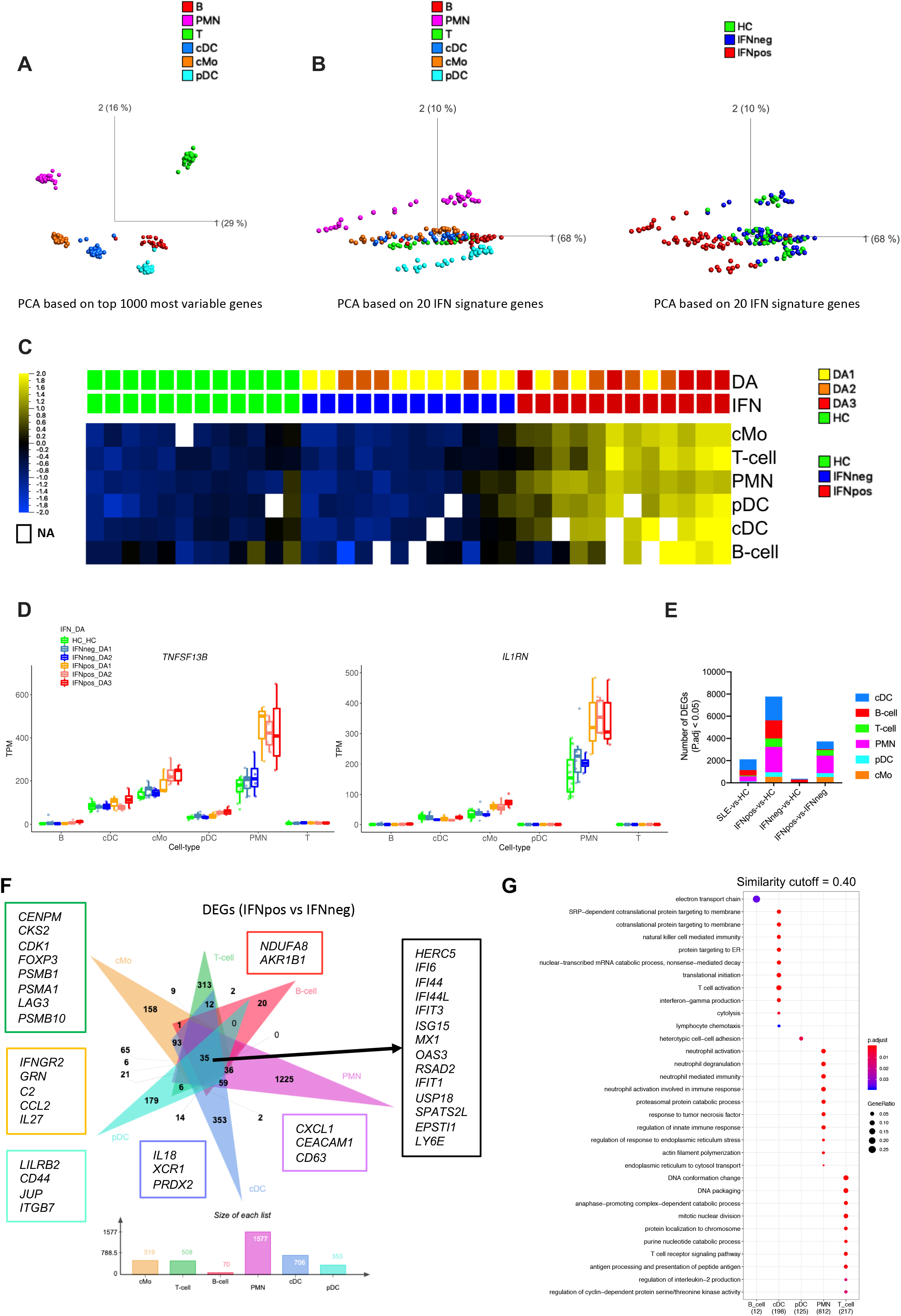
IFN-based molecular signature is independent of immune-cell types. **(A)** PCA plot based on 1000 most variable genes across six different immune cell types. The different cell types, B-cells, PMN, T-cells, cDC, cMo, and pDC, are shown in different colors, red, magenta, green, blue, orange, and cyan respectively. **(B)** PCA plots of different cell types based on IFN signature genes (IFN-20). The right plot is colored based on cell types and the left plot is colored based on IFN response status. **(C)** Average expression of IFN-20 genes presented as row-wise z-scores of TPM in IFNpos (red), IFNneg (blue) and HC (green) for each cell type separately. The upper panel shows SLEDAI score and is divided into three different categories based on score: DA1 (0-2), DA2 (3-7), and DA3 (>7) (yellow, orange and red, respectively). The detailed expression of each IFN-20 gene in all six cell types is provided (**Supplementary Figure S3G**). **(D)** Expression of *BAFF* and *IL1RN* across six different cell types using IFN response status and SLEDAI based categories. **(E)** A stacked column plot shows the number of RNA-Seq based differentially expressed genes with an adjusted P value of <0.05 (Benjamini-Hochberg test of DESeq2) in multiple cell type specific datasets and different comparisons. **(F)** The flower plot (generated by jvenn) shows the overlap of DEGs by IFNpos versus IFNneg (change with an adjusted P value of <0.05 (DESeq2 analysis; Benjamini-Hochberg test) in six different cell types. Some cell-specific genes of interest are highlighted in boxes (outline color is similar the color of each individual cell type). **(G)** Functional annotations (generated by clusterProfiler) of cellspecific DEGs from Figure 3F. We used similarity cutoff of 0.40 to remove similar gene-ontology terms. The color shows the significance (in terms of P.adj) and the size is gene ratio of annotations.

### Cell-specific transcriptional differences between two molecular subtypes of SLE

Even though the IFN response signature is shared and similar across all cell types, it is still essential to study each sorted immune cell type separately, which unlike bulk analysis of PBMCs, provides the resolution to identify cell type-specific transcriptional programs in relation to SLE pathogenesis. Therefore, we performed differential gene expression analysis comparing SLE-vs-HC, IFNpos-vs-HC, IFNneg-vs-HC, IFNpos-vs-IFNneg for each cell type separately (**Figure 3E; Supplementary Table S3**). We found only a few DEGs while comparing SLE-vs-HC whereas IFNpos-vs-HC comparison provided hundreds of DEGs in most of cell types (**Figure 3E**). We also observed a large number of DEGs between IFNpos and IFNneg groups for all cell types except B cells. B cells also have the largest number of DEGs for IFNneg-vs-HC analysis compared to all other cells and compared to IFNpos-vs-HC comparison in B cells. Most DEGs (P.adj < 0.05) are specific to one particular cell type and only 35 DEGs, all of which are a part of the IFN-363 set, are present in all cell types when we performed the IFNpos-vs-IFNneg comparison (**Figure 3F**). The gene-ontology based functional annotations of DEGs has revealed that all cell types were enriched for common IFN-related pathways (**Supplementary Figure S3H; Supplementary Table S4**).

The PMN, T cells, pDC, cDC, cMo, and B cells have 77.68% (1225/1577), 61.61% (313/508), 50.71% (179/353), 50.00% (353/706), 30.44% (158/519), and 28.57% (20/70) of their DEGs from the IFNpos vs IFNneg comparison only from that specific cell type, respectively (**Figure 3F**). These high number of cell type-specific DEGs suggest that, aside from the common IFN response genes, each cell type has different unique molecular changes between the two SLE subgroups. The functional enrichment analysis of these cell type-specific differences highlights several cell type-specific functions for each cell type (**Figure 3G; Supplementary Table S4**). The cDC-specific genes are involved in T cell activation (mainly *XCL1, IL18,* and *PRDX2* genes) through antigen presentation. Expression of *XCL1* (X-C Motif Chemokine Ligand 1) by crosspresenting CD8^+^ dendritic cells has been reported to determine cooperation with CD8+ T cells in murine models^48^. *IL18* is a pro-inflammatory cytokine that plays an important role in generating inflammation in lupus nephritis^49^. *PRDX2* (Peroxiredoxin 2) is an antioxidant that plays a key role in inflammation^50^. The pDC data are enriched for heterotypic cell-cell adhesion *(LILRB2, CD44, JUP, ITGB7)* and B cells have an enrichment of electron transport chain *(NDUFA8* and *AKR1B1)* related genes. PMN have genes enriched in neutrophil degranulation and neutrophil activation involved in immune response (*CXCL1*, *CEACAM1*, and *CD63*). T cells have enrichment differences in genes related to cell-cycle (*CENPM*, *CKS2*, *CDK1*), the T-receptor signaling pathway (*FOXP3*, *PSMB1*, *PSMA1*), and antigen processing and presentation of peptide antigen (*LAG3*, *PSMB10).* Even though we only mention a few genes here (**Figure 3F**), we also provide a complete list of genes and their GO-based functional enrichments for each cell type (**Figure 3G**; **Supplementary Figure S3H, Supplementary Table S4)**. The number of DEGs are relatively low for each cell type in our SLE-vs-HC comparisons (**Figure 3E**) highlighting the necessity of stratifying patient groups prior to comparative transcriptomic analysis.

### Integrated multi-cell type weighted gene co-expression analysis (iWGCNA)

It is accepted that multiple immune cells are involved in the complex SLE pathogenesis^23^ but how they really cross-talk with each other at the molecular level is largely unknown. The availability of multi-cell type transcriptome profiles from a sufficient number of patients provides us the unprecedented opportunity to develop and apply data-driven and integrative bioinformatics approaches to study this dataset. The standard WGCNA does not support multicell analysis; therefore, we developed a novel integrated WGCNA (iWGCNA) approach, which combines transcriptomes of different cell types by representing each gene with its name plus a cell type identifier to generate a single integrated weighted co-expression-based network. This integrated network resulted into 78 different modules from total 101,282 “genes” (nearly 17k genes from each cell type), with most of these modules containing genes from multiple cell types (**Figure 4A; Supplementary Table S5**). Such mixed modules suggest that genes from different cell types co-vary across patients and that a systematic quantification of such covariation could be useful for identifying molecular cross-talk among different cell types. Interestingly, most of the B cell genes are concentrated in one particular large module (turquoise) and have very little co-expression with other cell types, whereas genes from other cell types were distributed across many modules. In order to have differential expression analysis inform our interpretation of these modules, we also calculated the statistical significance of overlap with DEGs from each cell type for each module using a hypergeometric test. Out of the 78 modules, we find 13 with a significant enrichment of DEGs (IFNpos vs IFNneg) for at least one immune cell type and 4 (black, blue, lightyellow and darkolivegreen) where DEGs from at least two cell types are enriched significantly in the same module (**Figure 4B**).

**Figure 4.**
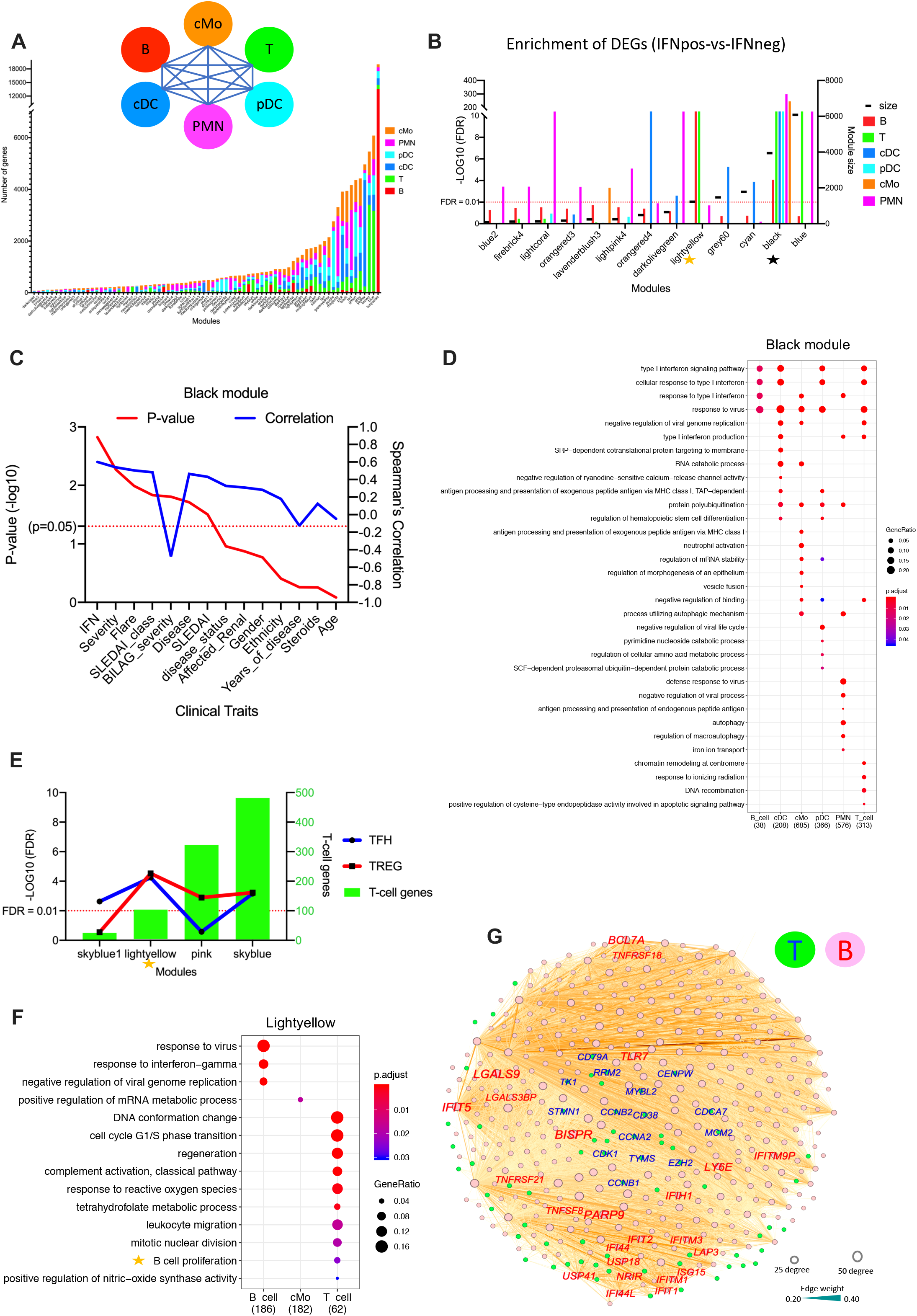
Integrated WGCNA (iWGCNA) analysis reveals IFN-driven cross-talk between T-cells and B-cells. **(A)** A figure depicting the iWGCNA analysis using combined transcriptomes from six different cell types. The iWGCNA has generated 78 different modules and a stacked column graph shows the proportion of genes from 6 different cell types in each module. The X-axis shows all modules and the Y-axis is the total number of genes in the corresponding modules. Gene numbers from different cell types are displayed in different colors as depicted in the cartoon. **(B)** We found 13 modules (out of total 78) that have significant enrichment of DEGs (IFNpos-vs-IFNneg) in at least one cell type. The X-axis shows modules with their enrichment in each cell type and Y1-axis shows −Log10 value of FDR (based on hypergeometric test). This includes four modules (black, blue, lightyellow, and darkolivegreen) where the DEGs are significantly enriched in two or more cell types. We have marked two modules (black and lightyellow) that we used for further analysis. Modules are sorted based on their increasing module size (showing as black dash on the Y2-axis). **(C)** The line plot shows significance (-Log10 of P-value) on Y1-axis in red) and Spearman’s correlation (Y2-axis in blue) of different clinical traits with Eigengene values of the black module from iWGCNA. The clinical traits are sorted (decreasing order) based on their statistical significance (-Log10 of P-value) for the black module. **(D)** Functional annotations (generated by clusterProfiler) of genes from each cell type in the black module. The color shows the significance (in terms of P.adj) and the size is gene ratio of annotations. **(E)** We have calculated enrichment of TFH and TREG gene sets in all 78 modules and found these four modules where they have significance for at least one gene set. The X-axis is different modules, the Y1-axis is -Log10 value of FDR (based on hypergeometric test), and the Y2-axis shows the number of genes from T-cells present in the corresponding module. Here also we marked the lightyellow module because only this module was also found in DEGs (IFNpos-vs-IFNneg) enrichment (**Figure 4B**) analysis. **(F)** The functional annotations of significantly enriched genes from different cell types in the lightyellow module. We highlighted the B cell proliferation function of T-cells because it is a known function of TFH. **(G)** T-cell and B-cell genes from the lightyellow module is visualized by Gephi. The nodes are colored according to cell of origin (T-cells in green and B-cells in red) and sized according to the number of edges (connections), and the edge thickness is proportional to the strength of co-expression. Genes from T-cells and B-cells are shown in blue and red, respectively.

### iWGCNA reveals IFN-driven cross-talk among different cell types

The black module we identified from iWGCNA consists of genes from each of the cell types and is relatively large with 3940 genes (1216 from cMo, 1004 from PMN, 594 from pDC, 534 from T cells, 520 from cDC, and 72 from B cells; **Supplementary Table S5**). Although, there are several clinical features such as severity, flare, SLEDAI, BILAG severity, which are significantly correlated with the black module, the correlation with IFN response status is the dominant feature (**Figure 4C-D; Supplementary Figure S4A**; **Supplementary Table S6**). Interestingly, the *BAFF* gene from three (cMo, PMN, and pDC) and the *IL1RN* gene from two different myeloid cell types (cMo and PMN) are also part of the black module showing co-expression with IFN-related genes across different immune cell types (**Supplementary Figure S4B**). The black module also harbors four genes coming from all six cell types including three that are IFN-related *(IFI6, IFIT3,* and *RSAD2*) and *ODF3B,* which is also likely regulated by *IRF3*^51^, a member of the interferon regulatory transcription factor (IRF) family. These results suggest that IFN response leads to coordination of gene expression changes across different cell types leading to a tightly connected iWGCNA module. Two other modules mentioned above (blue and darkolivegreen) (**Figure 4B**) show enrichment in harboring DEGs from two different cell types (IFNpos vs IFNneg comparison) but they lack clear functional enrichments for at least one of these two cell types (**Supplementary Figure S4C-D; Supplementary Table S6**).

### iWGCNA reveals IFN-driven cross-talk between T cells and B cells

Interestingly, the fourth module (lightyellow) mentioned above (**Figure 4B**) shows a significant enrichment for both B cell and T cell DEGs. This is particularly striking given that most B cell genes are readily clustered into one large module (turquoise) as discussed (**Figure 4A**). Since the natural connection between T and B cells is the T cell help for B cell function, and B cells are critical targets for SLE, we further looked into specific T helper subtypes that may correlate with enriched B cell function. The follicular helper T cells (TFH) are known to provide help and to play a central role in germinal center (GC) formation and development of high affinity antibodies and memory B cells^27^. Another T cell subset known as regulatory T cells (TREG) are also have been shown to directly suppress B-cells in SLE^52^ and *in vitro* in a TGF-beta dependent fashion^53^. Therefore, we computed the enrichment of published gene sets for TFH^26^ and TREG^41^ cells for each module identified from our iWGCNA analysis. To account for overlap among TFH, TREG and IFN response signatures, we removed genes from both the TFH and the TREG sets which overlap with any of the other two sets (**Supplementary Figure S4E**). Using the remaining 265 TFH and 300 TREG genes, we identified a total of four modules (lightyellow, pink, skyblue, and skyblue1) with significant overlap with at least one of these gene sets (**Figure 4E**) including the lightyellow module we identified earlier (**Figure 4B**). The functional annotation of this lightyellow module showed a significant enrichment of B cell proliferation-related genes (*CD38*, *MZB1,* and *CD79A)* expressed by T cells (**Figure 4F; Supplementary Table S6**). These B cell proliferation related T cell genes are also connected with other TFH and cell-cycle related genes from T cells. Most of thier co-expressed/connected genes from B cells are IFN response related but include members of the tumor necrosis factor (TNF) and tumor necrosis factor receptor (TNFR) superfamilies *(TNFSF8, TNFRSF18,* and *TNFRSF21)* and cross-talk related galectins (*LGALS9* and *LGALS3BP*), which are also well-connected in the lightyellow module (**Figure 4G**). We then revisited the multi-cell comparison, and found many genes that are characteristic of TFH cells (e.g., *TIGIT, STMN1, TYMS, FABP5, LAG3, CCNA2, CDKN3, CDCA7,* and *KPNA2),* and TREG cells (e.g., *FOXP3* and *HAVCR2/TIM3)* or are related to cell cycle (e.g., *CENPM, CKS2, PHF19, EZH2,* and *TK1*) and are also uniquely differentially expressed in IFNpos vs IFNneg comparisons for T cells only (**Figure 3F; Supplementary Table S3**).

The major advantage of iWGCNA is that it provides us a systematic way to combine gene expression and co-expression profiles across multiple cell types and results in a manageable number of modules or gene sets that then can be analyzed further with domain-specific knowledge. The identification of T cell and B cell cross-talk within a specific module in the context of SLE is a great example of iWGCNA’s power in teasing out such cross-cell type correlations with important biological and clinical implications.

### IFNpos patients have higher expression levels of genes related to TFH and TREG subsets

In order to better understand the influence of IFN response on composition of different T cell subsets, we use signature gene sets of TH1^54^, TH2^54^, TH17^54,55^, TFH^26^, and TREG^41^ from published studies (**Supplementary Table S7**). As before, we remove the IFN response related genes (IFN-363 set; **Supplementary Figure S4E**) from each T helper signature subset first. We then perform Gene Set Enrichment Analysis (GSEA) between IFNpos and IFNneg patients for each gene set. While, TFH (NES = 1.77, q = 0.006) and TREG (NES = 1.68, q = 0.009) gene sets show significant enrichment in IFNpos vs IFNneg comparison (**Figure 5A**), the other three subsets, TH1 (NES = 0.83, q = 0.694), TH2 (NES = 1.17, q = 0.326), and TH17 (NES = 1.47, q = 0.077), show no such enrichment (**Supplementary Figure S5A**).

**Figure 5.**
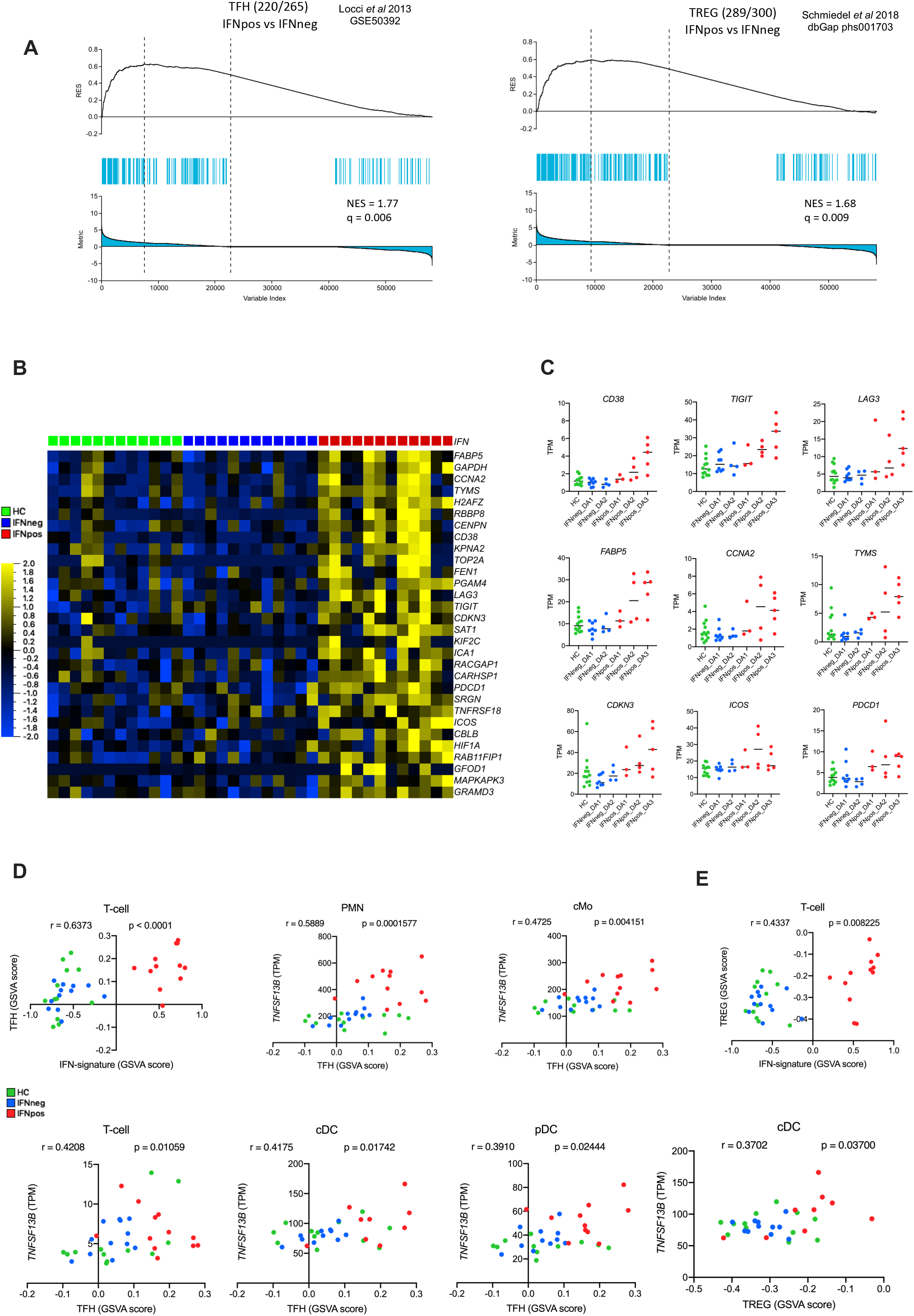
TFH and TREG signatures are more enriched in IFNpos patients and *BAFF* expression correlates with TFH feature from multiple cell types. **(A)** GSEA of TFH (left) and TREG (right) in the transcriptome of IFNpos versus IFNneg in T-cells, presented as running enrichment score (RES) for the gene set, from most over-represented genes on the left to the most underrepresented on the right; values above the plot represent the normalized enrichment score (NES) and false discovery rate (FDR)-corrected significance value; Kolmogorov-Smirnov test. The source of the gene set is also provided. **(B)** Expression of top 30 TFH-related genes, where each gene is presented as row-wise z-scores of TPM values in IFNpos (red), IFNneg (blue) and HC (green); each column represents an individual patient. **(C)** Individual expression plots for known TFH related genes divided by IFN response status as well as SLEDAI categories. **(D)** The top left plot shows Spearman’s correlation between GSVA score from IFN-20 genes (X-axis) and TFH gene set (Y-axis). Similarly, we calculated the correlation between GSVA score of TFH gene set and expression (TPM values) of *BAFF* (TNFSF13B) from different cell types. We found that *BAFF* expression in PMN, cMo, T-cells, cDC, and pDC has significant correlation with the TFH signature gene set (based on GSVA score). The significance (P-value) of correlation for all plots is also provided. **(E)** The top right plot shows Spearman’s correlation between GSVA score from IFN-20 genes (X-axis) and TREG gene set (Y-axis). We found significant correlation between *BAFF* expression and TREG signature gene set (based on GSVA score) from only cDC cells.

We then investigated the top TFH-related genes (based on filter by variance from statistics function of Qlucore) to find that mostly they have higher expression in IFNpos group whereas no or very little expression was observed in both the IFNneg and HC groups (**Figure 5B**). Many important TFH-related genes *(CD38, TIGIT, LAG3, FABP5, CCNA2, TYMS,* and *CDKN3)* are significantly differentially expressed between T cells from IFNpos vs IFNneg patients (**Figure 5C; Supplementary Table S3**). In general, expression levels of these genes are elevated in higher disease activity groups (DA2 and DA3) within IFNpos patients. Similarly, TREG-related genes including *FOXP3* and *HAVCR2/TIM3,* also have higher expression in the IFNpos group (**Supplementary Figure S5B**). These analyses show that both TFH-like and TREG-like molecular programs are enriched in IFNpos patients, though it is not clear whether the TREG-like signature is simply a response marker or plays a non-trivial role in SLE pathogenesis.

### *BAFF* expression from multiple cell types correlate with expression of TFH related genes

*BAFF* has previously been reported to regulate TFH cells and IFN-gamma production in lupus-prone mice^30^. Furthermore, increased expression of *BAFF* also induces expansion of activated B cells through TFH cells^31^. Our identification of *BAFF* from the combined DEGs and DCGs analysis of classical monocytes (**Figure 2D**), together with the significant enrichment of TFH related gene expression in T cells from IFNpos SLE patients suggest that there could be potential connection between *BAFF* expression and TFH-like features in the SLE patient blood. Since multiple immune cells produce *BAFF* (**Figure 3D**), it is also important to understand which cell types significantly contribute to the induction of TFH-like programs in T cells through the *BAFF* axis.

To this end, we employed the Gene Set Variation Analysis (GSVA), which estimates variation of enrichment of a particular gene set over a sample population^56^. GSVA provides enrichment scores for each sample for a given gene set, which can then be correlated with sample-specific features such as a gene expression measurement or another GSVA enrichment score. First, we found a strong positive correlation (r = 0.637, p < 0.0001) between GSVA scores from IFN-signature genes (IFN-20) and from TFH-related genes using expression profiles from T cells (**Figure 5D**). Next, we computed the correlation between *BAFF* expression from different immune cells and GSVA enrichment of TFH-related genes in T-cells resulting in significant correlations for all cells, except B cells, with specifically strong correlations for PMN (r = 0.589, p = 0.0001) and cMo (r = 0.4725, p = 0.004) (**Figure 5D; Supplementary Figure S5C)**. Similar analysis for TREG-related genes also highlighted some statistically significant correlations, though to a lesser extent compared to TFH-related genes (**Figure 5C, E; Supplementary Figure S5C)**. Although only correlative, these results suggest that *BAFF* production from PMN and cMo cells, concordant with increase in IFN response-related gene expression, may induce an extrafollicular T cell help program, akin to follicular B cell help by TFH cells, in IFNpos SLE patients. This in turn leads to a robust autoantibody production by B cells leading to formation of immune complexes and SLE manifestations. Further mechanistic studies are needed to completely characterize this interplay among IFN response by all immune cells, BAFF expression of PMN and cMo and enrichment of B cell activation by turning on specific T cell programs.

## DISCUSSION

The diagnostic complexity and heterogeneous nature of SLE disease require stratification of patients with the goal of correlating clinical features with molecular subgroups. Although, we have used variety of available clinical and demographic information (e.g., age, ethnicity, flare, severity, SLEDAI, and different treatments), IFN response status consistently provided the most striking relationships with gene expression differences and with gene co-expression modules (**Figure 2A; Figure 2B**). With this in mind, we analyzed SLE patients in two distinct groups, namely IFNpos and IFNneg throughout this work. The joint analysis of DEGs and DCGs between IFNpos and IFNneg in classical monocytes (**Figure 2C**) reveals two known immune modulators (*BAFF* and *IL1RN*) within a small group of prioritized genes, as well as several other genes of interest that could be further explored as therapeutic targets or biomarkers for SLE (**Figure 2D**). Although, IFN response status was mainly stable across different visits of the same patient, our cohort had one patient whose IFN response status changed between visits and this change was accompanied by changes in the serum levels and mRNA levels of pro-inflammatory cytokines (IL-6, IP-10/*CXCL10*, IFN-gamma; **Figure 1G**). Interestingly, the expression and co-expression profiles of *BAFF* and *IL1RN* also changed with IFN response status in this longitudinal data (**Figure 2E, H**). Recently, a phase III clinical trial (TULIP 2, NCT02446899) in SLE patients with a human monoclonal antibody (Anifrolumab) that specifically blocks *IFNAR1* (type I interferon receptor subunit 1) was completed^57^. The trial results also suggested that knowing patients’ IFN response status before prescribing targeted therapy will be helpful in the clinical setting and in designing further targeted therapies for specific patient groups.

Here we systematically analyzed the transcriptome profiles of multiple circulating immune cell types and found that IFN signature-based classification of SLE patients into two groups is very consistent across different immune cell types (**Figure 3C**). The large and clear-cut transcriptional differences between gene expression profiles of IFNpos and IFNneg patient groups across different immune cell types (**Figure 3F**) without such clear differences in IFN levels in the serum from the two groups, suggest that the cells might be pre-exposed to IFN signals prior to migrating into the blood. Beyond the shared IFN response signature, it has been challenging to infer biological functions of these cell-specific DEGs because GO-based functional annotations are generic and only provide information about canonical functions regardless of cell type. Recently, an integrated and multicohort analysis also has been published unified SLE MetaSignature of 93 genes in the blood and most of genes (85%) were IFN related^58^.

Another challenge was the involvement of multiple different immune cell types each of which added another layer of complexity. Here we proposed and demonstrated the use of a novel bioinformatics approach we named iWGCNA in exploring the cross-talk among different immune cell types using co-expression relationships of gene pairs across different cell types. We then revisited the large number of DEGs identified from one-by-one DEG analysis of IFNpos vs IFNneg patients **(Figure 3F**) using iWGCNA to explore potentially linked functional enrichments involving at least two different cell types. Out of 4 modules, which we narrowed down from a total of 78, one module corresponded mainly to the IFN response signature with an enrichment of IFNpos vs IFNneg DEGs from all six different cell types (**Figure 4B; Figure 4C**).

Interestingly, another module was significantly enriched for both T cell and B cell DEGs (**Figure 4B; Figure 4G**) and suggested that extrafollicular T cell help to B cells, i.e. gene signatures in the blood that are reminiscent of TFH cells, could play an important role in heightened disease activity in the IFNpos patient group (**Figure 4E; Figure 5A**). Recently, multiple lines of evidence have accumulated that circulating follicular helper-like T-cells are present in SLE blood and their expansion is associated with disease activity^59–61^. Another recent work also identified a T helper subset that is distinct from TFHs but still promotes B cell activation and differentiation into plasmablasts extrafollicularly in SLE blood^28^. It has been also shown that T peripheral helper (Tph) cells (PD-1^hi^CXCR5^-^CD4^+^) stimulates B cell responses in lupus via MAF and IL-21^62^. New evidences are also accumulating that T cell metabolism^63^ and epigenetic part^64^ also regulates the TFH cell in SLE. Our results do not exclude either possibility but highlight that such B cell help related T cell programs are likely more relevant targets for the treatment of IFNpos SLE patients, and not for IFNneg SLE patients. Furthermore, our finding that *BAFF* from three different myeloid cell types come together in the black module with significant co-expression to IFN-related genes (**Supplementary Figure S4B**), suggests that the effectiveness of a belimumab-based treatment could be impacted by the IFN response status of the patient.

In conclusion, we have found an IFN-based molecular heterogeneity in transcriptome profiles of SLE patients that is consistent across different immune cell types. We developed a novel computational approach (iWGCNA) to understand how this IFN-signal could affect the cross-talk among different immune cells. We anticipate that the data source generated here will be very useful for further characterization of immune cell type-specific roles in SLE progression. We also believe that our data-driven bioinformatics approach to jointly analyze gene expression profiles from multiple immune cell types from the same tissue (i.e., blood) will be broadly applicable to studies of other heterogeneous autoimmune diseases where multiple cell types are implicated in disease pathogenesis.

## AUTHOR CONTRIBUTIONS

B.P., P.V., and F.A. conceived the work, designed and analyzed experiments; B.P. performed RNAseq, iWGCNA, and data analysis under the supervision of P.V., and F.A. The bulk and single cell RNAseq experiments done by S.L., B.W., and G.S. The SLE cohort was conceived by R.S. and was generated by K.K., A.J.McK., R.S., and E.R. including patient recruitment, obtaining consent, disease activity assessment and sample collection. E.R. developed and performed the cell sorting strategy and flow cytometry analyses. B.P. wrote the first draft of manuscript that was revised and edited by P.V., F.A., A.J.McK., and R.S. All authors read and approved the final text of the manuscript.

## ACKNOWLEDGEMENTS

Interactive Fund grant from Kyowa Kirin Pharmaceutical Research, Inc. to P.V. The HiSeq2500 (Illumina) instrument was purchase by NIH S10OD016262 grant. We thank Anne-Laure Perraud for providing useful insights in this study.

## DATA AVAILABILITY

The accession number for the RNA sequencing data reported in this paper is NCBI GEO: GSE149050. All data and gene-specific expression pattern across different cell types is also available online through website: https://ay-lab-tools.lji.org/sle (username: reviewer, password: 695w6ZCwRH5Nwrye).

## SUPPLEMENTARY TABLES

**Supplementary Table S1.** The differential gene expression analysis result (DESeq2 output) between SLE and HC from first study visit data of classical monocytes (related to Figure 1B; Figure 1C).

**Supplementary Table S2.** The combined analysis results of differentially expressed genes (DEGs) and differentially connected genes (DEGs) between IFNpos and IFNneg in classical monocytes (related to Figure 2C).

**Supplementary Table S3.** The differential gene expression analysis results of multiple comparisons for six different immune cell types (related to Figure 3E; Figure 3F).

**Supplementary Table S4.** Details of gene-ontology based functional annotations of cell type specific DEGs between IFNpos and IFNneg (related to Figure 3G).

**Supplementary Table S5.** Comprehensive details of cell type, gene and corresponding module of iWGCNA based network (related to Figure 4A).

**Supplementary Table S6.** Details of gene-ontology based functional annotations of different cell type related genes of important iWGCNA modules (related to Figure 4D; Figure 4F).

**Supplementary Table S7.** Details of different (TFH, TREG, TH1, TH2, and TH17) gene sets that were used in the GSEA and GSVA analyses (related to Figure 5A).

## REFERENCES

1. Kaul, A. et al. Systemic lupus erythematosus. Nat. Rev. Dis. Prim. 2, 16039 (2016).

2. Davidson, A. What is damaging the kidney in lupus nephritis? Nat. Rev. Rheumatol. 12, 143–153 (2016).

3. Bayry, J. Lupus pathogenesis: role of IgE autoantibodies. Cell Res. 26, 271–272 (2016).

4. Carter, E. E., Barr, S. G. & Clarke, A. E. The global burden of SLE: prevalence, health disparities and socioeconomic impact. Nat. Rev. Rheumatol. 12, 605–620 (2016).

5. Obermoser, G. & Pascual, V. The interferon-alpha signature of systemic lupus erythematosus. Lupus 19, 1012–9 (2010).

6. Tsokos, G. C. Systemic Lupus Erythematosus. N. Engl. J. Med. 365, 2110–2121 (2011).

7. Lam, G. K. W. & Petri, M. Assessment of systemic lupus erythematosus. Clin. Exp. Rheumatol. 23, S120–32 (2005).

8. Bombardier, C., Gladman, D. D., Urowitz, M. B., Caron, D. & Chang, C. H. Derivation of the SLEDAI. A disease activity index for lupus patients. The Committee on Prognosis Studies in SLE. Arthritis Rheum. 35, 630–40 (1992).

9. Thanou, A., Chakravarty, E., James, J. A. & Merrill, J. T. Which outcome measures in SLE clinical trials best reflect medical judgment? Lupus Sci. Med. 1, e000005 (2014).

10. Touma, Z. & Gladman, D. D. Current and future therapies for SLE: obstacles and recommendations for the development of novel treatments. Lupus Sci. Med. 4, e000239 (2017).

11. Baechler, E. C. et al. Interferon-inducible gene expression signature in peripheral blood cells of patients with severe lupus. Proc. Natl. Acad. Sci. 100, 2610–2615 (2003).

12. Feng, X. et al. Association of increased interferon-inducible gene expression with disease activity and lupus nephritis in patients with systemic lupus erythematosus. Arthritis Rheum. 54, 2951–2962 (2006).

13. Crow, M. K. Type I interferon in the pathogenesis of lupus. J. Immunol. 192, 5459–68 (2014).

14. Der, E. et al. Single cell RNA sequencing to dissect the molecular heterogeneity in lupus nephritis. JCI Insight 2, (2017).

15. Chu, S. et al. The transcriptional program of sporulation in budding yeast. Science 282, 699–705 (1998).

16. de la Fuente, A. From ‘differential expression’ to ‘differential networking’ - identification of dysfunctional regulatory networks in diseases. Trends Genet. 26, 326–333 (2010).

17. Fuller, T. F. et al. Weighted gene coexpression network analysis strategies applied to mouse weight. Mamm. Genome 18, 463–472 (2007).

18. van Nas, A. et al. Elucidating the Role of Gonadal Hormones in Sexually Dimorphic Gene Coexpression Networks. Endocrinology 150, 1235–1249 (2009).

19. Southworth, L. K., Owen, A. B. & Kim, S. K. Aging Mice Show a Decreasing Correlation of Gene Expression within Genetic Modules. PLoS Genet. 5, e1000776 (2009).

20. Oldham, M. C., Horvath, S. & Geschwind, D. H. Conservation and evolution of gene coexpression networks in human and chimpanzee brains. Proc. Natl. Acad. Sci. U. S. A. 103, 17973–8 (2006).

21. Stark, R., Grzelak, M. & Hadfield, J. RNA sequencing: the teenage years. Nature Reviews Genetics 20, 631–656 (2019).

22. Tsokos, G. C., Lo, M. S., Reis, P. C. & Sullivan, K. E. New insights into the immunopathogenesis of systemic lupus erythematosus. Nat. Rev. Rheumatol. 12, 716–730 (2016).

23. Moulton, V. R. et al. Pathogenesis of Human Systemic Lupus Erythematosus: A Cellular Perspective. Trends Mol. Med. 23, 615–635 (2017).

24. Langfelder, P. & Horvath, S. WGCNA: an R package for weighted correlation network analysis. BMC Bioinformatics 9, 559 (2008).

25. Simpson, N. et al. Expansion of circulating T cells resembling follicular helper T cells is a fixed phenotype that identifies a subset of severe systemic lupus erythematosus. Arthritis Rheum. 62, 234–244 (2010).

26. Locci, M. et al. Human circulating PD-1+CXCR3-CXCR5+ memory Tfh cells are highly functional and correlate with broadly neutralizing HIV antibody responses. Immunity 39, 758–769 (2013).

27. Crotty, S. T Follicular Helper Cell Differentiation, Function, and Roles in Disease. Immunity 41, 529–542 (2014).

28. Caielli, S. et al. A CD4 + T cell population expanded in lupus blood provides B cell help through interleukin-10 and succinate. Nature Medicine 25, 75–81 (2019).

29. Mackay, F. & Browning, J. L. BAFF: A fundamental survival factor for B cells. Nature Reviews Immunology 2, 465–475 (2002).

30. Coquery, C. M. et al. BAFF regulates follicular helper T cells and affects their accumulation and interferon-γ production in autoimmunity. Arthritis Rheumatol. 67, 773–784 (2015).

31. Chen, M. et al. The function of BAFF on T helper cells in autoimmunity. Cytokine and Growth Factor Reviews 25, 301–305 (2014).

32. Subramanian, A. et al. Gene set enrichment analysis: A knowledge-based approach for interpreting genome-wide expression profiles. Proc. Natl. Acad. Sci. U. S. A. 102, 15545–15550 (2005).

33. Zhang, J. et al. Type I interferon related genes are common genes on the early stage after vaccination by meta-analysis of microarray data. Hum. Vaccines Immunother. 11, 739–745 (2015).

34. Banchereau, R. et al. Personalized Immunomonitoring Uncovers Molecular Networks that Stratify Lupus Patients. Cell 165, 551–565 (2016).

35. Rönnblom, L. & Leonard, D. Interferon pathway in SLE: One key to unlocking the mystery of the disease. Lupus Sci. Med. 6, (2019).

36. Lai, K. N. & Yap, D. Y. H. Cytokines and their roles in the pathogenesis of systemic lupus erythematosus: From basics to recent advances. Journal of Biomedicine and Biotechnology 2010, (2010).

37. Cui, Y. X., Fu, C. W., Jiang, F., Ye, L. X. & Meng, W. Association of the interleukin-6 polymorphisms with systemic lupus erythematosus: A meta-analysis. Lupus 24, 1308–1317 (2015).

38. Kong, K. O. et al. Enhanced expression of interferon-inducible protein-10 correlates with disease activity and clinical manifestations in systemic lupus erythematosus. Clin. Exp. Immunol. 156, 134–140 (2009).

39. Furst, D. E. Anakinra: Review of recombinant human interleukin-I receptor antagonist in the treatment of rheumatoid arthritis. Clin. Ther. 26, 1960–1975 (2004).

40. Sims, J. E. & Smith, D. E. The IL-1 family: regulators of immunity. Nat. Rev. Immunol. 10, 89–102 (2010).

41. Schmiedel, B. J. et al. Impact of Genetic Polymorphisms on Human Immune Cell Gene Expression. Cell 175, 1701–1715.e16 (2018).

42. Wegiel, B. & Otterbein, L. E. Go Green: The Anti-Inflammatory Effects of Biliverdin Reductase. Front. Pharmacol. 3, 47 (2012).

43. Takaoka, A. et al. DAI (DLM-1/ZBP1) is a cytosolic DNA sensor and an activator of innate immune response. Nature 448, 501–505 (2007).

44. Kuriakose, T. & Kanneganti, T. D. ZBP1: Innate Sensor Regulating Cell Death and Inflammation. Trends in Immunology 39, 123–134 (2018).

45. Grünvogel, O. et al. DDX60L Is an Interferon-Stimulated Gene Product Restricting Hepatitis C Virus Replication in Cell Culture. J. Virol. 89, 10548–68 (2015).

46. Wang, X. et al. The Cellular Antiviral Protein APOBEC3G Interacts with HIV-1 Reverse Transcriptase and Inhibits Its Function during Viral Replication. J. Virol. 86, 3777–3786 (2012).

47. Liu, H., Luo, K. & Luo, D. Guanosine monophosphate reductase 1 is a potential therapeutic target for Alzheimer’s disease. Sci. Rep. 8, 2759 (2018).

48. Dorner, B. G. et al. Selective Expression of the Chemokine Receptor XCR1 on Crosspresenting Dendritic Cells Determines Cooperation with CD8+ T Cells. Immunity 31, 823–833 (2009).

49. Mohsen, M. A., Abdel Karim, S. A., Abbas, T. M. & Amin, M. Serum interleukin-18 levels in patients with systemic lupus erythematosus: Relation with disease activity and lupus nephritis. Egypt. Rheumatol. 35, 45–51 (2013).

50. Knoops, B., Argyropoulou, V., Becker, S., Ferté, L. & Kuznetsova, O. Multiple roles of peroxiredoxins in inflammation. Molecules and Cells 39, 60–64 (2016).

51. Rouillard, A. D. et al. The harmonizome: a collection of processed datasets gathered to serve and mine knowledge about genes and proteins. Database 2016, baw100 (2016).

52. Iikuni, N., Lourenço, E. V., Hahn, B. H. & La Cava, A. Cutting Edge: Regulatory T Cells Directly Suppress B Cells in Systemic Lupus Erythematosus. J. Immunol. 183, 1518–1522 (2009).

53. Xu, A. et al. TGF-β-Induced Regulatory T Cells Directly Suppress B Cell Responses through a Noncytotoxic Mechanism. J. Immunol. 196, 3631–41 (2016).

54. Arlehamn, C. L. et al. Transcriptional Profile of Tuberculosis Antigen-Specific T Cells Reveals Novel Multifunctional Features. J. Immunol. 193, 2931–2940 (2014).

55. Hu, D. et al. Transcriptional signature of human pro-inflammatory TH17 cells identifies reduced IL10 gene expression in multiple sclerosis. Nat. Commun. 8, (2017).

56. Hänzelmann, S., Castelo, R. & Guinney, J. GSVA: gene set variation analysis for microarray and RNA-Seq data. BMC Bioinformatics 14, 7 (2013).

57. Morand, E. F. et al. Trial of Anifrolumab in Active Systemic Lupus Erythematosus. N. Engl. J. Med. 382, 211–221 (2020).

58. Haynes, W. A. et al. Integrated, multicohort analysis reveals unified signature of systemic lupus erythematosus. JCI Insight 5, (2020).

59. Choi, J. Y. et al. Circulating follicular helper-like T cells in systemic lupus erythematosus: Association with disease activity. Arthritis Rheumatol. 67, 988–999 (2015).

60. Xu, H. et al. Increased frequency of circulating follicular helper T cells in lupus patients is associated with autoantibody production in a CD40L-dependent manner. Cell. Immunol. 295, 46–51 (2015).

61. Zhang, X. et al. Circulating CXCR5+CD4+helper T cells in systemic lupus erythematosus patients share phenotypic properties with germinal center follicular helper T cells and promote antibody production. Lupus 24, 909–917 (2015).

62. Bocharnikov, A. V. et al. PD-1hiCXCR5-T peripheral helper cells promote B cell responses in lupus via MAF and IL-21. JCI Insight 4, (2019).

63. Sharabi, A. & Tsokos, G. C. T cell metabolism: new insights in systemic lupus erythematosus pathogenesis and therapy. Nature Reviews Rheumatology 16, 100–112 (2020).

64. Su, C. et al. Human follicular helper T cell promoter connectomes reveal novel genes and regulatory elements at SLE GWAS loci. bioRxiv 2019.12.20.885426 (2019). doi:10.1101/2019.12.20.885426

