## Supplementary Information for "Integrative transcriptomic analysis of SLE reveals IFN-driven cross-talk between immune cells"

### SUPPLEMENTARY FIGURES

#### Figure S1

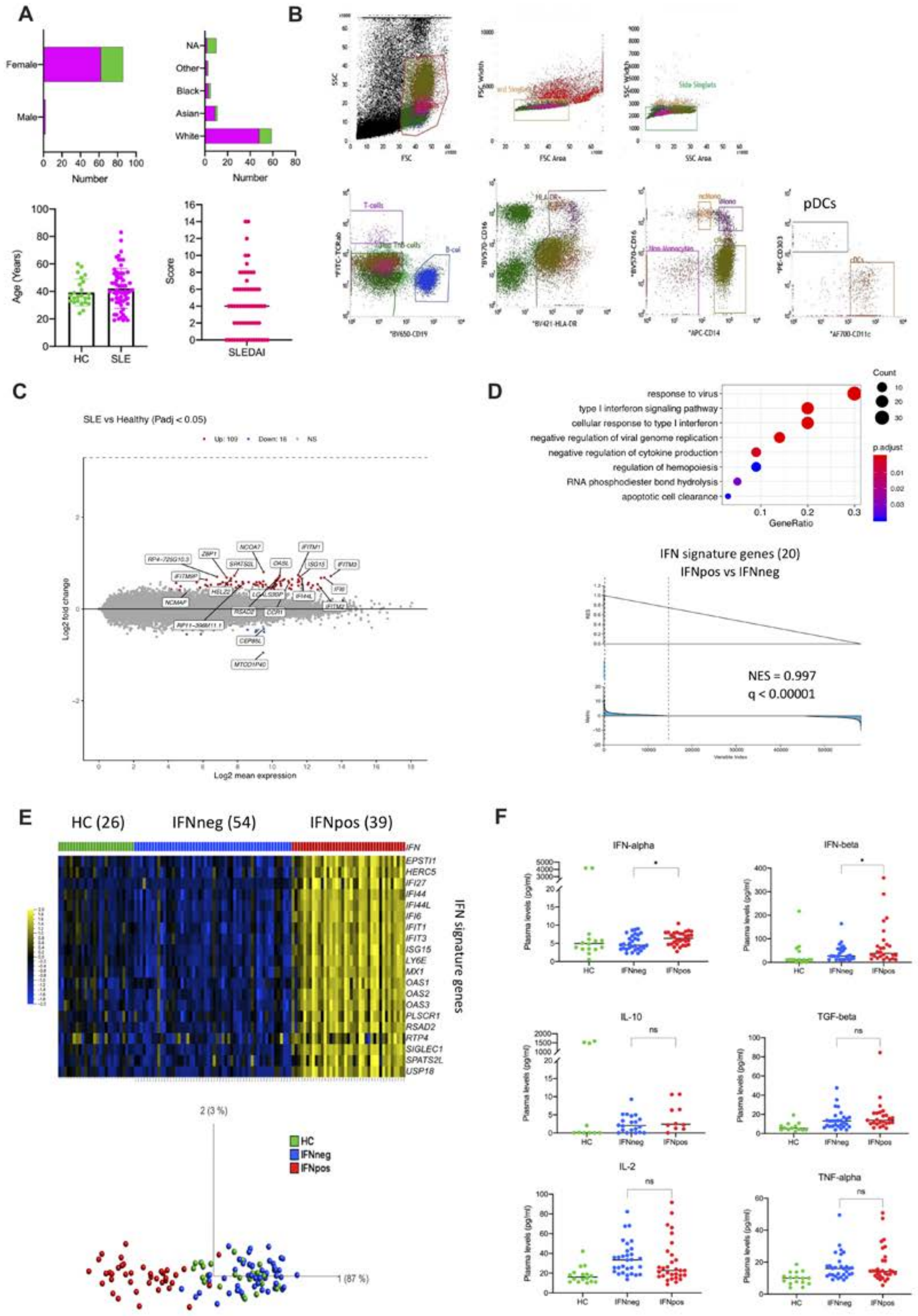

**Supplementary Figure S1. Transcription profile of classical monocytes reveals two molecular subtypes of SLE that shows stability in longitudinal data. (A)** Different plots show the distribution of gender, ethnicity, age, and SLEDAI score in the SLE cohort. Healthy control (HC) samples are highlighted in green color and SLE samples are highlighted in magenta color. **(B)** Gating scheme of T-depleted PBMC for sorting of multiple immune cell types. Live cells were gated based on their forward side scatter versus side scatter profile followed by single cells gating by FSC then SSC and then finally according to their cell surface phenotype as follows, B cells (CD19+, TCRa/b-), classical Mo (cMono, TCRa/b-, CD19-, HLA-DR+,CD14+, CD16-), intermediate Mo (iMono, TCRa/b-, CD19-, HLA-DR+, CD16+, CD14+), non-classical Mo (ncMono, TCRa/b-, CD19-, HLA-DR+, CD16+ CD14 low), plasmacytoid dendritic cells (pDC, TCRa/b-, CD19-, HLA-DR+, CD303+, CD11c-) and conventional dendritic cells (cDC, TCRa/b-, CD19-, HLA-DR+, CD303-, CD11c+). **(C)** The MAplot shows on 125 DEGs (P.adj < 0.05 from Benjamini-Hochberg test in DESeq2) between SLE and HC in classical monocytes. The top 20 genes (based on P.adj) are highlighted in boxes. **(D)** Functional annotations (generated by clusterProfiler) of DEGs between IFNpos-vs-IFNneg (top). The color shows the significance (in terms of P.adj), the size is gene counts in annotation, and the X-axis shows gene ratio. The GSEA plot shows significant enrichment (NES=0.997; q<0.00001) of 20 IFN-signature genes (IFN-20) in IFNpos vs IFNneg comparison. **(E)** All SLE patients (including longitudinal visits) classified into two groups based on the expression of IFN-20 genes (top). Where each gene is presented as row-wise z-scores of transcripts per million (TPM) in IFNpos (red), IFNneg (blue) and HC (green); each column represents an individual patient. The PCA plot (bottom) shows the two molecular sub-types of SLE in different colors. Green, blue, and red represent HC, IFNneg and IFNpos, respectively (we used this color scheme in all figures). **(F)** Scatter dot-plots show plasma level expression (pg/ml) of different cytokines. Only first visit samples (n=76) were used in these scatter dot-plots. Differences between IFNpos and IFNneg were calculated using unpaired T-test (two-tailed) and statistical significance (p-value) levels are shown in each plot (ns: not significant, *: <0.05).

### Figure S2

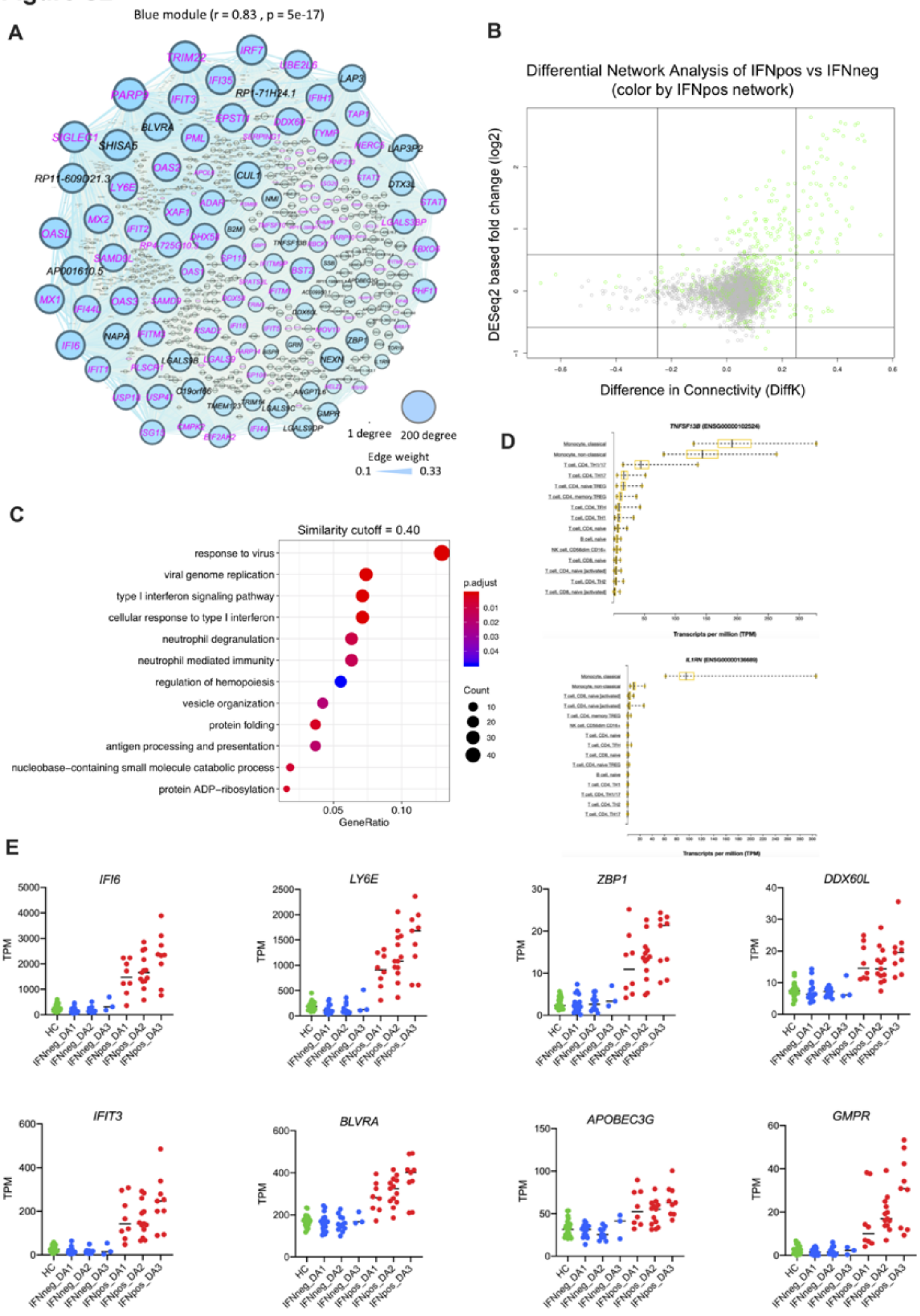
A
Blue module (r = 0.83 , p = 5e-17)
1 degree
200 degree
Edge weight
0.1
0.33
B
Differential Network Analysis of IFNpos vs IFNneg
(color by IFNpos network)
DESeq2 based fold change (log2)
Difference in Connectivity (DiffK)
C
Similarity cutoff = 0.40
response to virus
viral genome replication
type I interferon signaling pathway
cellular response to type I interferon
neutrophil degranulation
neutrophil mediated immunity
regulation of hemopoiesis
vesicle organization
protein folding
antigen processing and presentation
nucleobase−containing small molecule catabolic process
protein ADP−ribosylation
p.adjust
Count
GeneRatio
D
E
IFI6
LY6E
ZBP1
DDX60L
IFIT3
BLVRA
APOBEC3G
GMPR
TPM
HC
IFNneg_DA1
IFNneg_DA2
IFNneg_DA3
IFNpos_DA1
IFNpos_DA2
IFNpos_DA3

**Supplementary Figure S2. Combined analysis of differential network and gene expression of classical monocytes reveals two known immune modulators (*BAFF* and *IL1RN*) and whose expression is dysregulated in SLE. (A)** Gephi based visualization of the blue module, where nodes are sized according to the number of edges (connections), and the edge thickness is proportional to the strength of co-expression. Available IFN related genes (IFN-363) are highlighted in magenta colors. **(B)** The plot shows both differentially expressed (DEGs) and connected genes (DCGs) from the IFNpos network based green and grey modules only. Where the X-axis is the difference in the connectivity (DiffK = K1-K2; K1=connectivity in IFNpos network; K2=connectivity in IFNneg network) and the Y-axis is the DESeq2 based fold-change (log2). By default, the 'grey' module is generated by WGCNA for non-co-expressed genes so it shows that green module genes are well-expressed as well as well-connected in the IFNpos in comparison to grey module genes. **(C)** Functional annotations (generated by clusterProfiler) from all green module genes in IFNpos network. The color shows the significance (in terms of P.adj), the size is gene counts in annotation, and the X-axis shows gene ratio. **(D)** The DICE database-based expression of *BAFF* and *IL1RN* in different immune cell types. Where X-axis is expression (TPM) level and Y-axis shows different immune cell types. **(E)** The individual expression plot of genes of interest from the green module using IFN response status as well as SLEDAI categories.

Figure S3

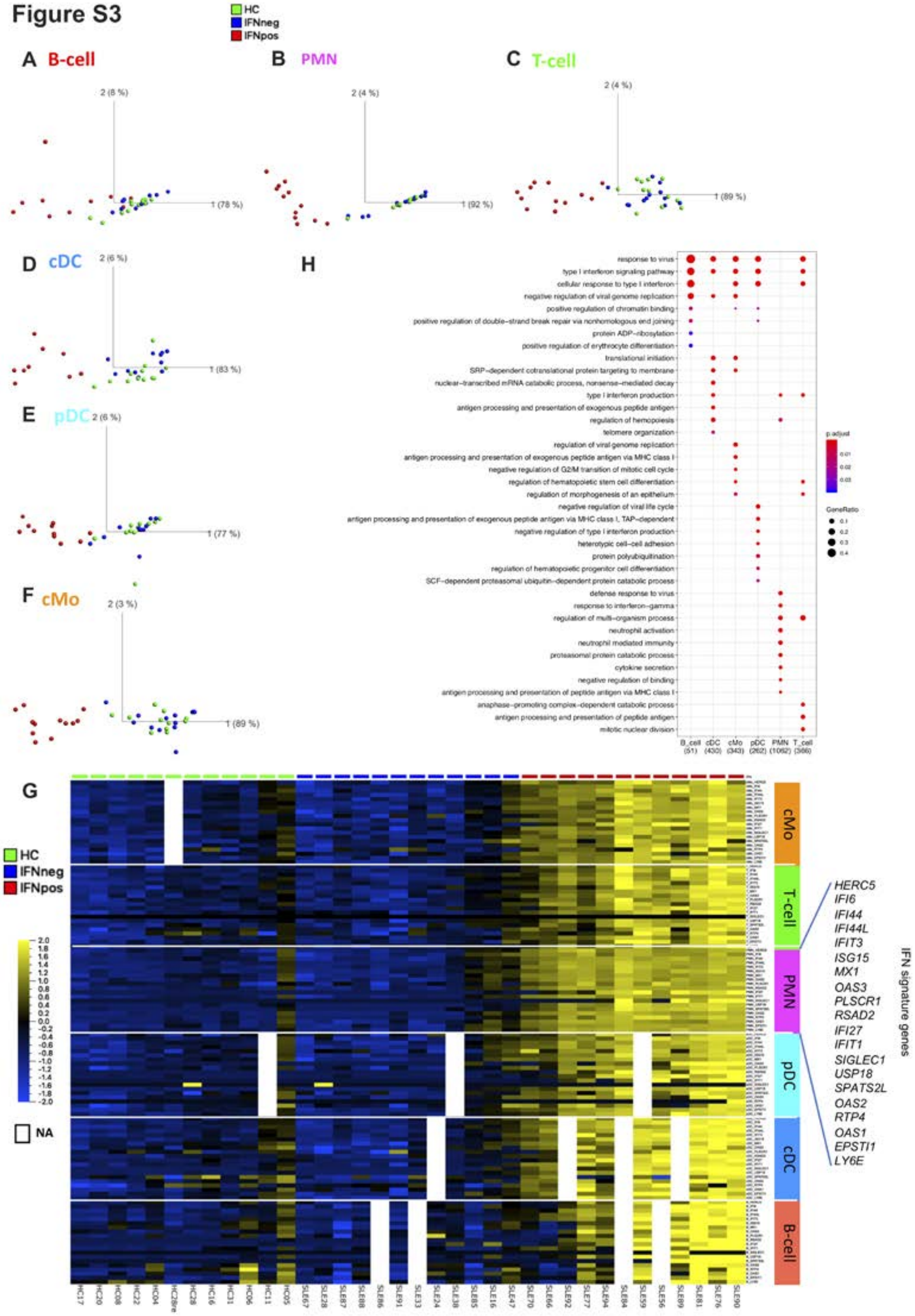


**Supplementary Figure S3. IFN-based molecular signature is independent of immune cell types. (A-F)** PCA plots of different cell types based on IFN signature genes (IFN-20). **(G)** Expression of IFN-20 genes presented as row-wise z-scores of TPM in IFNpos (red), IFNneg (blue) and HC (green) for each cell type separately. The expression profile is absent (used blank or ‘NA’) for a few patients in some cell types. **(H)** Functional annotations (generated by clusterProfiler) of all DEGs (IFNpos-vs-IFNneg) from different cell types. The color shows the significance (in terms of P.adj) and the size is gene ratio of annotations.

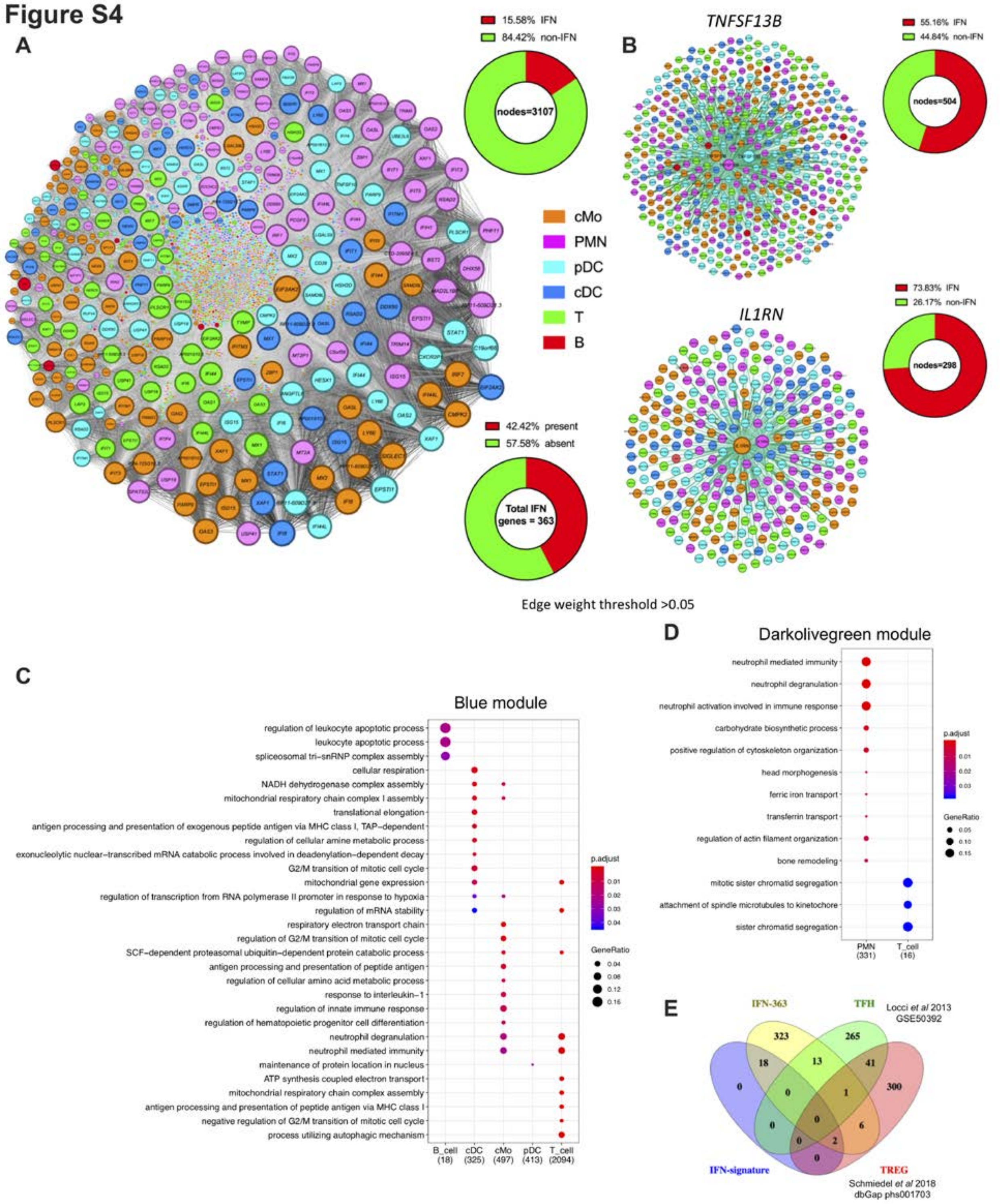


**Supplementary Figure S4. Integrated WGCNA (iWGCNA) analysis reveal IFN-driven cross-talk between T-cells and B-cells. (A)** The network of the black module genes visualized by Gephi.

The nodes are colored according to cell of origin and sized according to the number of edges (connections), and the edge thickness is proportional to the strength of co-expression. The top donut chart shows that 15.58% nodes are IFN-related genes (from IFN-363) and the bottom donut plot shows that 42.42% of IFN-363 genes, from at least one-cell type, are present in the black module. **(B)** The Gephi based visualization of *BAFF* (top) and *IL1RN* (bottom) connected genes in the black module and the corresponding donut chart show the proportion of IFN and non-IFN related nodes (based on IFN-363 genes). *BAFF* from three different cell types (cMo, PMN, and pDC) and *IL1RN* from cMo and PMN are present in the black module. **(C-D)** Functional annotations (generated by clusterProfiler) of genes from each cell type in the blue **(C)** and darkolivegreen **(D)** modules. The color shows the significance (in terms of P.adj) and the size is gene ratio of annotations. **(E)** A Venn diagram shows the overlap of TFH and TREG signature genes with IFN-20 and IFN-363 genes. We only used genes that are exclusively present in TFH (265) and TREG (300) signatures.

### Figure S5

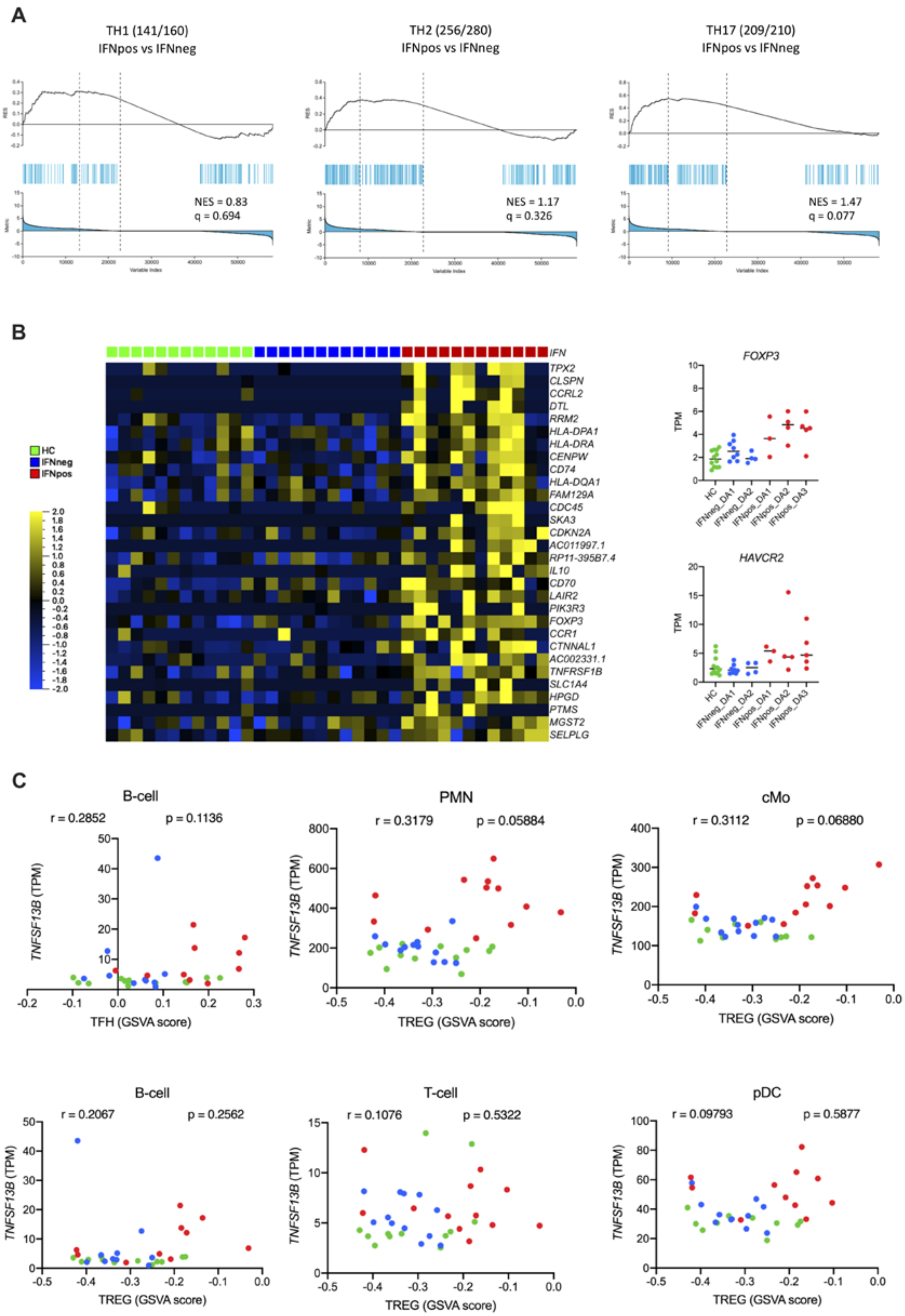


**Supplementary Figure S5. TFH and TREG signatures are more enriched in IFNpos patients and *BAFF* expression in multiple cell types correlates with TFH feature. (A)** GSEA of TH1 (left), TH2

(middle), and TH17 (right) in the transcriptome of IFNpos versus IFNneg in T-cells, presented as running enrichment score (RES) for the gene set, from most over-represented genes at left to most under-represented at right; values above the plot represent the normalized enrichment score (NES) and false discovery rate (FDR)-corrected significance value; Kolmogorov-Smirnov test. **(B)** A heatmap (left) shows expression of the top 30 TREG-related genes, where each gene is presented as row-wise z-scores of TPM values in IFNpos (red), IFNneg (blue) and HC (green); each column represents an individual patient. Individual expression plots (right) of two TREG related genes (*FOXP3* and *HAVCR2*) using IFN response status as well as SLEDAI categories. **(C)** The correlation between GSVA score of TFH and TREG gene set and expression (TPM values) of *BAFF* from different cell types. The Spearman's correlation with significance value (p-value) is given.

### METHODS

#### Patient samples

Healthy volunteers were recruited by the Normal Donor Blood programs at La Jolla Institute for Immunology (La Jolla, CA) and The Scripps Research Institute (La Jolla, CA). SLE patients were recruited by Division of Rheumatology, University of California, San Diego and Sanguine Bioscience. All subjects recruited for this study followed Institutional Review Boards (La Jolla Institute for Immunology, La Jolla, CA, University of California San Diego, La Jolla, CA, The Scripps Research Institute, La Jolla, CA, and Quorum Review, Seattle, WA) approvals and study participants gave written informed consent. SLE Patients were previously diagnosed by a clinician and classified as active if they had a SLEDAI score of at least 4 as well as a BILAG score of A or B at each study visit. A subset of patient samples (n=10) were collected by Sanguine Bioscience and patient information collected via self-reporting of active symptoms. Longitudinal samples were collected from 17 SLE patients and 2 healthy individuals from September 2014 through August 2016 for 1 to 5 follow-up visits (**Figure 1G**).

#### Sample collection and cell isolation

After the volunteers consented to donate their blood sample, 40 mL of venous blood was withdrawn from their arms with a needle. 36 mL were collected for sorting and flow cytometric analysis (6 x 6 mL BD vacutainer tubes with potassium EDTA (Becton Dickinson, Franklin Lakes, NJ) and 4 mL for serum analysis. Whole blood was spun at 2000 rpm for 10 minutes. The plasma fraction was transferred to a fresh tube, aliquotted and stored at -80C until analyzed. The cell pellets were then resuspended in HBSS without calcium and magnesium and underlayed with Ficoll-paque Plus (Fisher Scientific). The tubes were centrifuged at 2000 rpm at room temperature for 20 minutes in sealed carriers with the brake off. The top fraction (plasma) was removed and frozen for future analysis. The interface layer containing the peripheral blood mononuclear cells (PBMC) was collected using a sterile rubber bulb and Pasteur pipette and transferred into a new 50 mL conical tube. The collected PBMC were washed in PBS by centrifuge at 1800 rpm, at room temperature for 10 minutes. The PBMC were passed through a 70 μm cell strainer, washed again and then either frozen in 10% Fetal Bovine Serum (GE

Healthcare Life Sciences) in DMSO or processed for purification of immune cells subsets by cell sorting as described below.

The cell pellets from the Ficoll gradients were used for isolation of PMN by density gradient centrifugation. First Red Blood Cells were lysed by resuspending the cell pellet in Gey's solution. Following incubation on ice for 5 min cells the solution was underlayed with 5 mL FBS and then centrifuged at 1800 rpm for 5 minutes. The cells were resuspended in PBS, passed through a 70 μm cell strainer and washed twice with PBS by centrifugation at 1200 rpm for 5 minutes each. PMN were resuspended in 30 mL HBSS without calcium or magnesium. 6 mL of 6% Dextran was added to the cells and mixed by inversion. Following incubation at room temperature for 30 minutes the supernatant was transferred to a new tube which was spun for 5 minutes at 1200 rpm. The pellet containing the PMN was then resuspended with 5 mL $H_2O$ and mixed for 5 seconds before addition of HBSS to a final volume of 50 mL. After a final spin at 1200 rpm for 5 minutes the PMN were resuspended in 5 mL HBSS, counted, spun again at 1200 rpm for 5 minutes. 1 mL of TRIZOL LS (Thermo Fisher) was added for each 5 x $10^6$ cells and the PMN samples were stored at -80C.

**Cell sorting**

Total T cells were isolated from PBMC by magnetic bead-based separation, specifically using CD3 DynaBeads (Thermo Fisher) according the manufacturer's instructions. The purified T cells were resuspended in Trizol LS and stored at -80C. The T-depleted PBMC were resuspended in FACS buffer (PBS, 2% FBS, 2 mM EDTA and 25 mM HEPES) containing mouse IgG (Jackson Immunochemicals) to block non-specific binding. Following incubation for 10 minutes on ice the cells were stained with a cocktail of antibodies to TCRαβ, CD11c, HLA-DR, CD16, CD19 (Biolegend, San Diego, CA), CD14 (Becton Dickinson, Mountainview, CA) and CD303 (Miltenyi, Bergisch Gladbach, Germany) **(Supplementary Figure 1B)**. B cells, classical, non-classical, and intermediate monocytes, pDC, and cDC were sorted on either a FACSAria or a BD Infux (Becton Dickinson). After sorting the cells were washed, lysed in Trizol LS and frozen at -80C immediately. Circulating classical $CD14^{hi}$ CD16- Monocytes were isolated from PBMC from 64 patients with Systemic Lupus Erythematosus (SLE) and 24 healthy subjects (**Figure 1A**). Five additional immune cell types (B-cells, T-cells, cDC, pDC, and PMN) were evaluated from a subset of the samples (24 SLE and 12 HC).

**Cytokine profiling of plasma**

Plasma was collected from patients. Frozen plasma was shipped to Affymetrix (Vienna, Austria) and analyzed in a 34-protein vendor-defined multiplex Procarta Plex-2panel (ThermoFisher, Santa Clara, CA) to profile differential plasma protein expression from healthy volunteers and patients with SLE. Analytes measured included soluble CD40 ligand, CXCL5, IFN-α2, IFN-β, IFN-γ, IL-10, IL-23, IL-12p70, IL-15, IL-17A, IL-17F, IL-18, IL-1α, IL-1b, IL-1RA, IL-2, IL-21, IL-4, IL-6, IP-10, CXCL11, MCP-1, RANTES, TGFβ TNFα, VEGF-A, IL-13, Leptin, PAI-1, Resistin, Fas ligand, SDF-1, IL-22, and GM-CSF. GraphPad Prism8 was used to generate scatter plots and to perform statistical analyses of these data.

**Bulk RNA sequencing**

Total RNA was isolated from sorted cell populations using miRNAeasy Micro kit (Qiagen, USA) and quantified. 3ng of total RNA was used to generate cDNA following the Smart-seq2 protocol. cDNA was purified using AMPure XP beads (0.8x, Beckman Coulter). Next, 1ng of cDNA from each sample was used to generate a sequencing library (Nextera XT DNA sample preparation kit and index kit, Illumina). The libraries were pooled and sequenced on a HiSeq2500 (Illumina) to obtain 50-bp single end reads. Both full-transcriptome amplification and sequencing library preparations were performed in a 96-well format to reduce assay-to-assay variability. Quality control steps were included after each step to eliminate samples with low quality from downstream process. A detailed protocol has been previously published [1]. Libraries were sequenced on a HiSeq2500 Illumina to obtain a minimum of 10 million 50-bp single-end reads (HiSeq SR Cluster Kit v4 cBot, HiSeq SBS Kit v4).

**Single-cell sequencing**

Cells from three SLE (IFNpos) patients were sorted into plates to generate sequencing libraries following the SmartSeq2 protocol. cDNAs were purified twice with AmPure XT beads (0.8x). 0.4ng of cDNA from each sample was used to generate a sequencing library using the Nextera XT DNA sample preparation kit and index kit from Illumina. Quality control steps were included after each step to eliminate samples with low quality from downstream process. A total of 156-single cell libraries passed all quality control criteria. The libraries were pooled then sequenced on a HiSeq 2500 to obtain a minimum of 50 thousand 50-bp single-end reads. A detailed protocol has been previously published [1].

**RNA-Seq analysis**

Bulk RNA-seq data (FASTQ files) were mapped against the hg38 genome (GRCh38.p7) reference using TopHat [2]; v2.0.9 (--max-multihits 1 --microexon-search --bowtie1) with FastQC (v0.11.2), and Samtools v0.1.19.0 [3]. Trimmomatic (v0.36) was used to remove adapters [4]. We employed htseq-count -m union -s no -t exon -i gene_name (part of the HTSeq framework, version v0.7.1 [5]) for calculating read counts. To identify differentially expressed genes between two groups, we used raw read counts and performed negative binomial tests for unpaired comparisons using the DESeq2 (v1.14.1) with package from Bioconductor [6]. We have disabled the default options of DESeq2 for independent filtering and Cooks cutoff. Non-expressed (no reads in all samples) genes have been filtered out before running DESeq2. All genes with Benjamini-Hochberg–adjusted P value of < 0.05 (based on DESeq2 results) have considered as differentially expressed genes (DEGs) in any comparison. The MAplot was generated by using ggmaplot function of ggpubr R package. Gene expression values were normalized as transcripts per million (TPM) and applied in the Qlucore Omics Explorer 3.3 software package for visualization and representation (heat maps, principal component analysis, and GSEA) of RNA-Seq data. Top-30 TFH/TREG genes were selected based on filter by variance from statistics function of Qlucore. Different box-plots and Spearman's correlation plots were generated by GraphPad Prism8 (v8.3.0).

**Weighted Gene Co-expression Network Analysis (WGCNA)**

R package WGCNA (v1.61) was used to generate co-expression network from the TPM data matrix [7]. To develop standard WGCNA network for SLE (**Figure 2A**), we used 16444 well expressed genes with TPM >1 in at least 25% of the samples and modules were generated using blockwiseModules function (parameters: checkMissingData = TRUE, power = 5, TOMType = unsigned, minModuleSize = 30, maxBlockSize = 16444, mergeCutHeight = 0.25). The pickSoftThreshold function was used to optimize soft-thresholding power (β) by choosing the lowest power for which the scale-free topology fit index reaches 0.90. The default 'grey' module generated by WGCNA for non-co-expressed genes. As each module by definition is comprised of highly correlated genes, their combined expression may be usefully summarized by eigengene profiles, effectively the first principal component of a given module. A small number of eigengene profiles may therefore effectively 'summarize' the principle patterns within the cellular transcriptome with minimal loss of information. This dimensionality-reduction approach also facilitates correlation of ME with traits. Different clinical features (IFN-status, age, ethnicity, flare, severity, SLEDAI score, years of disease, affected renal, BILAG severity, and different treatments) were used as a trait and correlated with MEs. Significance of correlation between this trait and MEs was assessed using Spearman's correlation and p-values.

In the differential network analysis (**Figure 2C**), we used standard WGCNA-based approach to generate co-expression networks for IFNpos and IFNneg separately. For each network, genes were clustered into a dendrogram and modules were assigned by blockwiseModules function (parameters: checkMissingData =TRUE, power = 6, TOMType = unsigned, minModuleSize = 30, maxBlockSize = 16444, mergeCutHeight = 0.25). In order to make both IFNpos and IFNneg networks comparable, we used same soft-thresholding power (β=6). Furthermore, network connectivity values using softConnectivity function (power=6) have been calculated for each gene where connectivity (also known as degree) is defined as the sum of connection strengths (based on co-expression) with the other genes in the network. The difference between the connectivity (DiffK = K1 – K2) for each gene between IFNpos (K1) and IFNneg (K2) was calculated (as described in [8]). Genes with at least ±0.25 difference (K1-K2) in connectivity (DiffK) were considered as differentially connected genes (DCGs).

**Integrated Weighted Gene Co-expression Network Analysis (iWGCNA)**
In the integrated WGCNA (iWGCNA) approach (**Figure 4A**), we have generated a single WGCNA network by merging transcriptome profiles of six different cell types (patient-matched) together. We used a total of 25 samples (10 HC, 8 IFNneg, and 7 IFNpos) from six different cell types. Highly correlated genes from combined transcriptomes across six immune cell types were identified and a total of 78 modules were generated using blockwiseModules function (parameters: checkMissingData = TRUE, power = 6, TOMType = unsigned, minModuleSize = 50, maxBlockSize = 101282, mergeCutHeight = 0.40). In order to find gene set specific important modules, we have measured the significance of a particular gene set (e.g. DEGs between IFNpos and IFNneg) by hypergeometric test using phyper R function and further p-values were adjusted for multiple test correction using p.adjust R function (method=fdr).

To visualize co-expression networks, we used the function exportNetworkToCytoscape at weighted = true, threshold = 0.05. A soft thresholding power was chosen based on the criterion of approximate scale-free topology. Networks were generated in Gephi (v0.9.2) using Fruchterman Reingold and Noverlap functions [9]. The size and color were scaled according to the average degree as calculated in Gephi, while the edge width was scaled according to the WGCNA edge weight value.

**Gene Set Enrichment Analysis (GSEA) and Gene Set Variation Analysis (GSVA)**
GSEA determines whether an a priori defined 'set' of genes (such as a signature) show significant cumulative changes in gene expression between phenotypic subgroups [10]. We applied GSEA using Qlucore Omics Explorer 3.3 software package for assessing significant enrichment of specific gene sets (eg. IFN signatures or T cell subtypes) in one group relative to that in another group (eg. IFNpos versus IFNneg). In summary, first all genes are ranked on the basis of their differential expression (TPM-based) in one group versus their expression in another group. Thereafter, a running enrichment score (RES) is calculated for a provided gene set on the basis of how often its genes appear at the top or bottom of the already ranked differential list. A default of 1,000 random permutations of the phenotypic subgroups are used to establish a null distribution of RES against which a normalized running enrichment score (NES) and false-discovery-rate-corrected q values are calculated using Kolmogorov-Smirnov statistic. We ran GSEA with different gene sets of TH1 [11], TH2 [11], TH17 [11,12], TFH [13], and TREG [14] from published studies (**Supplementary Table S7**) after removing IFN-20 and IFN-363 related genes (**Supplementary Figure S4E**) to uncover only T-cell subtypes specific enrichments. These gene signatures were selected to test the null hypothesis that IFN based sub-groups (IFNpos and IFNneg) did not show significant enrichment for different T cell sub-types.

In order to establish correlation between two different gene sets or groups, we need to calculate enrichment score for each sample. The GSVA [15] estimates variation of enrichment of particular gene set over a sample population and provides enrichment score for each sample. GSVA was implemented using gsva function of R package GSVA (v1.20.0) with rnaseq=TRUE parameter and it provided GSVA scores that we used to correlate different gene sets with *BAFF* expression (**Figure 1D; Figure 1E; Supplementary Figure S5C**).

**Gene-Ontology (GO) based functional annotations**
The biological relevance of important genes from different analyses was further investigated using the clusterProfiler [16]. To functionally annotate genes (eg. DEGs or module genes) from one cell type, we used enrichGO function (parameters: OrgDb = org.Hs.eg.db, ont = BP). We also removed redundant GO-terms (parameters: cutoff=0.40, by=p.adjust, select_fun = min, measure = Wang) with more than 40% similarity cut-off. In the plots (**Supplementary Figure S1D; Supplementary Figure S2C**), color shows the significance (in terms of P.adj), size is gene counts in annotation, and X-axis shows gene ratio. To compare multiple cell types, compareCluster function (parameters: fun = enrichGO, OrgDb = org.Hs.eg.db, ont = BP) was used to generate Gene-Ontology based comparative functional annotations of different cell types. We used all available genes from different cell types in a module to run clusterProfiler and only displayed cell types that have significant enrichment of any GO-term. In multi-cell plots (**Figure 3G; Figure 4D; Figure 4F**), color displays the significance (in terms of P.adj) of particular GO-terms and size shows the gene ratio of annotations.

**Quantification and statistical analysis**

Statistical analyses were performed using GraphPad Prism8 (v8.3.0). The Spearman's correlation coefficient (r value) was used to access the significance of correlations between the levels of any two components of interest. R packages were applied with R version 3.3.3 using x86_64-pc-linux-gnu (64-bit) platform under CentOS Linux 7 (Core).

**REFERENCES**


1. Rosales, S. L. *et al.* A Sensitive and Integrated Approach to Profile Messenger RNA from Samples with Low Cell Numbers. *Methods Mol. Biol.* **1799**, 275–302 (2018).
2. Trapnell, C., Pachter, L. & Salzberg, S. L. TopHat: discovering splice junctions with RNA-Seq. *Bioinformatics* **25**, 1105–11 (2009).
3. Li, H. *et al.* The Sequence Alignment/Map format and SAMtools. *Bioinformatics* **25**, 2078–2079 (2009).
4. Bolger, A. M., Lohse, M. & Usadel, B. Trimmomatic: A flexible trimmer for Illumina sequence data. *Bioinformatics* **30**, 2114–2120 (2014).
5. Anders, S., Pyl, P. T. & Huber, W. HTSeq-A Python framework to work with high-throughput sequencing data. *Bioinformatics* **31**, 166–169 (2015).
6. Love, M. I., Huber, W. & Anders, S. Moderated estimation of fold change and dispersion for RNA-seq data with DESeq2. *Genome Biol.* **15**, 550 (2014).
7. Langfelder, P. & Horvath, S. WGCNA: an R package for weighted correlation network analysis. *BMC Bioinformatics* **9**, 559 (2008).
8. Fuller, T. F. *et al.* Weighted gene coexpression network analysis strategies applied to mouse weight. *Mamm. Genome* **18**, 463–472 (2007).
9. Clarke, J. *et al.* Single-cell transcriptomic analysis of tissue-resident memory T cells in human lung cancer. *J. Exp. Med.* **216**, 2128–2149 (2019).
10. Subramanian, A. *et al.* Gene set enrichment analysis: A knowledge-based approach for interpreting genome-wide expression profiles. *Proc. Natl. Acad. Sci. U. S. A.* **102**, 15545–15550 (2005).
11. Arlehamn, C. L. *et al.* Transcriptional Profile of Tuberculosis Antigen–Specific T Cells Reveals Novel Multifunctional Features. *J. Immunol.* **193**, 2931–2940 (2014).
12. Hu, D. *et al.* Transcriptional signature of human pro-inflammatory TH17 cells identifies reduced IL10 gene expression in multiple sclerosis. *Nat. Commun.* **8**, (2017).
13. Locci, M. *et al.* Human circulating PD-1+CXCR3-CXCR5+ memory Tfh cells are highly functional and correlate with broadly neutralizing HIV antibody responses. *Immunity* **39**, 758–769 (2013).
14. Schmiedel, B. J. *et al.* Impact of Genetic Polymorphisms on Human Immune Cell Gene Expression. *Cell* **175**, 1701-1715.e16 (2018).
15. Hänzelmann, S., Castelo, R. & Guinney, J. GSVA: gene set variation analysis for microarray and RNA-Seq data. *BMC Bioinformatics* **14**, 7 (2013).
16. Yu, G., Wang, L. G., Han, Y. & He, Q. Y. ClusterProfiler: An R package for comparing biological themes among gene clusters. *Omi. A J. Integr. Biol.* **16**, 284–287 (2012).
